# Targeted mutagenesis using CRISPR-Cas9 in the chelicerate herbivore *Tetranychus urticae*

**DOI:** 10.1101/2019.12.25.888032

**Authors:** Wannes Dermauw, Wim Jonckheere, Maria Riga, Ioannis Livadaras, John Vontas, Thomas Van Leeuwen

## Abstract

The use of CRISPR-Cas9 has revolutionized functional genetic work in many organisms, including more and more insect species. However, successful gene editing or genetic transformation has not yet been reported for chelicerates, the second largest group of terrestrial animals. Within this group, some mite and tick species are economically very important for agriculture and human health, and the availability of a gene-editing tool would be a significant advancement for the field. Here, we report on the use of CRISPR-Cas9 to create gene knock-outs in the spider mite *Tetranychus urticae*. The ovary of virgin adult females was injected with a mix of Cas9 and sgRNAs targeting the phytoene desaturase gene. Natural mutants of this gene have previously shown an easy-to-score albino phenotype. Albino sons of injected virgin females were mated with wild-type females, and two independent transformed lines where created and further characterized. Albinism inherited as a recessive monogenic trait. Sequencing of the complete target-gene of both lines revealed two different lesions at expected locations near the PAM site in the target-gene. Both lines did not genetically complement each other in dedicated crosses, nor when crossed to a reference albino line with a known genetic defect in the same gene. In conclusion, two independent mutagenesis events were induced in the spider mite *T. urticae* using CRISPR-Cas9, providing an impetus for genetic transformation in chelicerates and paving the way for functional studies using CRISPR-Cas9 in *T. urticae*.

## 1 Introduction

Mites and ticks are members of the chelicerates, the largest group of terrestrial animals after insects. *T. urticae* and other spider mites are important crop pests worldwide. This herbivore species is at the extreme end of the generalist-to-specialist spectrum and can feed on a staggering 1,100 plant species. Not surprisingly, it is currently reported as the ‘most resistant’ pest worldwide, as it developed resistance to more than 90 acaricides (Mota-Sanchez and Wise, 2019; Van Leeuwen and Dermauw, 2016; Van Leeuwen et al., 2015). In 2011, a 90 Mb high-quality Sanger-sequenced genome became available for this species (Grbic et al., 2011). This allowed to disentangle some of the molecular mechanisms underlying resistance, whether to man-made pesticides or plant secondary compounds. The extreme adaptation potential of *T. urticae* was associated with specific gene expansions in known detoxification enzyme families, such as cytochrome P450 monooxygenases, glutathione-S-transferases, carboxyl-choline esterases, an unexpected repertoire of ABC and MFS transporters, and a proliferation of cysteine peptidases (Dermauw et al., 2013a; Dermauw et al., 2013b; Grbic et al., 2011; Santamaría et al., 2012). In addition, several genes acquired via horizontal gene transfer were uncovered and characterized, such as intradiol-ring cleaving dioxygenases (Schlachter et al., 2019; Wybouw et al., 2012; Wybouw et al., 2014; Wybouw et al., 2018). Gene-expression studies have revealed large transcriptional differences between susceptible and resistant *T. urticae* strains, as well as after short-term transfer or adaptation to new hosts (Dermauw et al., 2013b; Grbic et al., 2011; Snoeck et al., 2018; Wybouw et al., 2014; Wybouw et al., 2015; Zhurov et al., 2014). Furthermore, mite-plant interactions have been thoroughly examined (Alba et al., 2015; Blaazer et al., 2018; Bui et al., 2018; Jonckheere et al., 2016; Martel et al., 2015; Santamaría et al., 2019; Wybouw et al., 2015; Zhurov et al., 2014). For instance, some salivary proteins were shown to modulate plant defenses (Iida et al., 2019; Villarroel et al., 2016). The availability of a high-quality genome and new technical advances in high-throughput sequencing has also led to the development of a genetic mapping tool, bulked-segregant analysis, which allowed to map quantitative trait loci at high resolution (Bryon et al., 2017; Kurlovs et al., 2019; Van Leeuwen et al., 2012; Wybouw et al., 2019a). To conclude, the spider mite has been an exceptional good model to study adaptation, owing to clear advantages in experimental manipulation, a small high-quality genome and the development of advanced genomic mapping tools.

However, the lack of tools for reverse genetics that can directly validate the involvement of genes and mutations in phenotypes of interest (and validate most of the work outlined above) has impeded critical advances in *T. urticae* molecular biology. RNA interference (RNAi) has dramatically accelerated scientific progress in different groups of insects (Scott et al., 2013), linking genes with phenotypes, but this technique is currently not always straightforward in mites (Kwon et al., 2016; Suzuki et al., 2017). Even more so, a recent technique, named <u>c</u>lustered <u>r</u>egularly <u>i</u>nterspaced <u>s</u>hort <u>p</u>alindromic <u>r</u>epeats (CRISPR) - <u>C</u>RISPR-<u>a</u>ssociated <u>p</u>rotein 9 (Cas9), has revolutionized functional genetic work in many organisms (Zhang and Reed, 2017). Successful CRISPR-Cas9-mediated gene manipulation has been reported for a steadily increasing number of organisms in the arthropod subphyla Crustaceae (Gui et al., 2016; Martin et al., 2016; Nakanishi et al., 2014) and Hexapoda, including Diptera, Hymenoptera, Hemiptera, Coleoptera, Orthoptera and diverse Lepidoptera (see Sun et al. (2017) for a review, Kotwica-Rolinska et al. (2019); Xue et al. (2018)), but not in the wide group of chelicerates. It is clear that the development of such method for directed, heritable gene editing is also crucial for the study of *T. urticae* and other mite and tick species.

The CRISPR-Cas9 technique currently usually consists of a two-component system with a small, easy to synthesize single guide RNA (sgRNA) and a bacterial nuclease (Cas9). It introduces double-stranded breaks in eukaryotic genomes, where the breaks can be repaired randomly (non-homologous end-joining, NHEJ) or based on a template (homology-directed repair) (Scott et al., 2018). In order to obtain efficient genomic DNA cleavage, Cas9 and sgRNA should be introduced into the oocytes (Rungger et al., 2017). In *Drosophila*, this is currently most easily accomplished by injecting sgRNAs in transgenic embryos expressing Cas9 under a germline-specific promotor (see for example Bajda et al. (2017) and Douris et al. (2016), and references in Korona et al. (2017)). Most current approaches with non-model organisms rely upon delivering the Cas9 ribonucleoprotein (RNP) complex (Cas9 protein + sgRNA) by embryonic microinjection (Chaverra-Rodriguez et al., 2018). However, within the chelicerates, embryo injection has not been accomplished yet, as injected embryos die (Garb et al., 2018; Sharma, 2017). This is probably the main reason why transgenic mites and ticks have not yet been reported (with the exception of one older study that was never replicated (Presnail and Hoy, 1992)). An alternative method, avoiding the injection of eggs or embryos, is delivery of the RNP complex to the germline by injecting the mother animals. Such approaches already proved to be successful for organisms such as nematodes (Witte et al., 2015) and insects (Chaverra-Rodriguez et al., 2018; Hunter et al., 2018; Macias et al., 2019). In this study, we used a similar approach, and injected virgin *T. urticae* females with a Cas9-sgRNA complex targeting the *T. urticae* phytoene desaturase gene, a gene essential for red pigmentation (Bryon et al., 2017; Bryon et al., 2013). Among the progeny, we identified albino males and show that their albino phenotype was the result of Cas9-induced mutations in the phytoene desaturase gene, hereby providing proof of principle of the feasibility of genetic modification of mites and other chelicerates.

## 2 Material and Methods

### 2.1 *T. urticae* strain

The London strain (wild type, WT) of *T. urticae* is an outbred reference laboratory strain (Van Leeuwen et al., 2012) and was used for sequencing of the complete *T. urticae* genome (Grbic et al., 2011). All injection experiments were performed with mites from this strain. The Alb-NL strain used in complementation tests was previously described (Bryon et al., 2017). All strains were maintained as previously described (Riga et al., 2017) on *Phaseolus vulgaris* cv. “Prelude” at 26±1°C, 60% RH and 16:8 (light:dark) photoperiod.

### 2.2 Recombinant Cas9 ribonucleoproteins and sgRNAs

Recombinant *Streptococcus pyogenes* Cas9 protein with an N-terminal nuclear localization signal (NLS) (Alt-R^®^ S.p. Cas9 Nuclease V3, catalog # 1081058) was purchased from Integrated DNA Technologies (Leuven, Belgium). Two guide sequences were designed using the CRISPOR website ((2018), accessed in December 2018), with the following settings: *T. urticae* phytoene desaturase sequence (*tetur01g11270*, https://bioinformatics.psb.ugent.be/orcae/overview/Tetur) as target (“Step 1”), *T. urticae* London genome (GCA_000239435.1) as genome (“Step 2”) and “20 bp NGG – Sp Cas9” as Protospacer Adjacent Motif (“Step 3”). Based on the guide DNA sequences, 3 nmol of single guide RNAs (sgRNA) was ordered. The ordered sgRNAs were synthetic sgRNAs (sgRNA1 and sgRNA2) from Synthego (Synthego Corporation, Menlo Park, California, USA), consisting of a 20 nt guide sequence (g1 or g2) + 80-mer “Synthego scaffold”

### 2.3 *In vitro* Cas9-sgRNA cleavage experiment

Before performing *in vivo* CRISPR-Cas9 experiments with *T. urticae*, we tested whether the Cas9-sgRNA complex could cleave PCR products of *tetur01g11270 in vitro*. Primer3 (Rozen and Skaletsky, 2000) was used to design primers that amplify the *tetur01g11270* regions that are targeted by the two sgRNAs (see above). An 895 bp region is amplified by the “tetur01g11270_DNA_1” primers (amplicon 1, containing the sgRNA1 cutting site), while “tetur01g11270_DNA_2” primers amplify a 699 bp region (amplicon 2, containing the sgRNA2 cutting site). *T. urticae* DNA was extracted from the WT strain using the Gentra Puregene Tissue Kit (QIAgen), according to the manufacturer’s instructions and using 100 adult females as starting material. The PCR of *tetur01g11270* fragments (amplicon 1 and 2) was conducted using the Expand™ Long Range dNTPack (Sigma-Aldrich). PCR reaction mixtures were prepared according to the manufacturer’s instructions and using the following temperature profile: denaturation for 2 min at 92°C, followed by five touch-down cycles of denaturation at 92°C for 10 s, annealing at 60°C −1°C/cycle for 15 s and elongation at 68°C for 1 min. Next, 37 cycles of 92°C for 10 s, 55°C for 15 s and 68°C for 1 min. After a final elongation of 68°C for 5 min, PCR products were checked by agarose gel electrophoresis, and purified using the EZNA^®^ Cycle Pure Kit (Omega Bio-Tek). The *in vitro* digestion protocol was performed as described by the IDT Alt-R CRISPR-Cas9 System Protocol (version September 2019, available at https://eu.idtdna.com/pages/support/guides-and-protocols, document ID# CRS-10096-PR 09/19), with some modifications. Briefly, the RNP complex was created by combining 2.5 µl sgRNA (10 µM stock in TE buffer, pH 7.5), 0.4 µl Alt-R S.p. Cas9 enzyme (62 µM stock) and 22.1 µl Cas9 dilution buffer (30 mM HEPES, 150 mM KCl, pH 7.5). For negative controls, sgRNA was replaced by TE. After incubation for 10 min at RT, the *in vitro* digestion reaction was assembled at RT as follows: 2 µl 10x Cas9 Nuclease Reaction Buffer (200 mM HEPES, 1 M NaCl, 50 mM MgCl_2_, 1 mM EDTA, pH 6.5), 4 µl Cas9 RNP (from previous step), 10 µl DNA substrate (amplicon 1 or 2, 50 nM stock) and 4 µl of water. The reaction mixture was incubated for 90 min at 37°C, after which 2 µL proteinase K (Sigma-Aldrich; 10 mg/ml) was added, and the DNA substrate was released from the Cas9 endonuclease by incubating for 10 min at 56°C. Subsequently, the digestion was analyzed using gel electrophoresis, in which 15 µL reaction mixture was loaded on gel.

### 2.4 *In vivo* Cas9-sgRNA cleavage experiment

#### 2.4.1 Cas9-sgRNA injection mix

The Cas9-sgRNA injection mix was prepared as indicated in Table S1. The final concentration of the Cas9 protein in the injection mix was 4.85 μg/μL (29.61 μM). Stock solution of each sgRNA was prepared by dissolving 3 nmol of sgRNA into 30 μL of RNAse-free water. sgRNAs were added to the injection mix in a 1:3 Cas9:sgRNA molar ratio and 0.5 mM of chloroquine was also included in the injection mix. The Cas9-sgRNA injection mix was incubated at 37°C for 10 min, and finally, the injection mix was centrifuged at 4°C for 10 min at 10,000 g and kept on ice until injection.

#### 2.4.2 Injection of *T. urticae* female mites

Female mites of the WT strain were allowed to lay eggs on the upper part of bean leaves on wet cotton in a Petri dish. After eight days, teliochrysalis females were transferred to another leaf disk and allowed to molt. After another one to four days, these unfertilized females were used for injections. Agar plates were made by dissolving 15 g of agar into 500 mL of cherry juice (for color contrast, brand “Eviva”) and subsequently heated until boiling. An agar “platform” was made by adding two glass microscope slides (26 × 76 mm, 1.1 mm thick; APTACA, Canelli, Italy), attached to each other by double-sided tape, into a Petri dish immediately after pouring the agar plates. After solidification of the agar, the microscope slides were removed, and the agar plate was cut in two along the length of the microscope slide (Figure S1). Unfertilized females were aligned on the agar platform, with their dorsal and right lateral side in contact with the agar (Figure 1). Injection needles were pulled from Clark capillary glass (borosilicate with filament: 1.0 mm (outside diameter, OD) × 0.58 (inner diameter, ID) × 100 mm (length); catalog # W3 30-0019/GC100F-10 (Harvard Apparatus Ltd, Holliston, Massachussets, USA)) using a P97-micropipette needle puller (Sutter Instruments, Novato, California, USA), with the following settings “Heat: 510, Pull: 20, Velocity: 90, Time: 250” (Figure S2). Mites were injected under a Leitz BIOMED Microscope (Wild Leitz/Leica, Wetzlar, Germany) and with a mechanical micromanipulator (Leitz/Leica, Wetzlar, Germany) that holds the injection needle (Figure 1). Approximately 6 nl of Cas9-sgRNA injection mix was injected in the ovary, near the third pair of legs, using a IM 300 Microinjector (Narishige, London, UK). Two batches (A and B) of mites were injected. Each batch of injected mites was transferred to a separate leaf disk and allowed to lay eggs. After 24 hours, the injected females were transferred to a new leaf disk and allowed to lay eggs again. The male haploid progeny of injected females (on six leaf disks in total (2 batches: A and B, 2 time-points: 0-24h and 24-48h)) was visually screened for the albino phenotype beginning 3 days after egg deposition.

**Figure 1.**
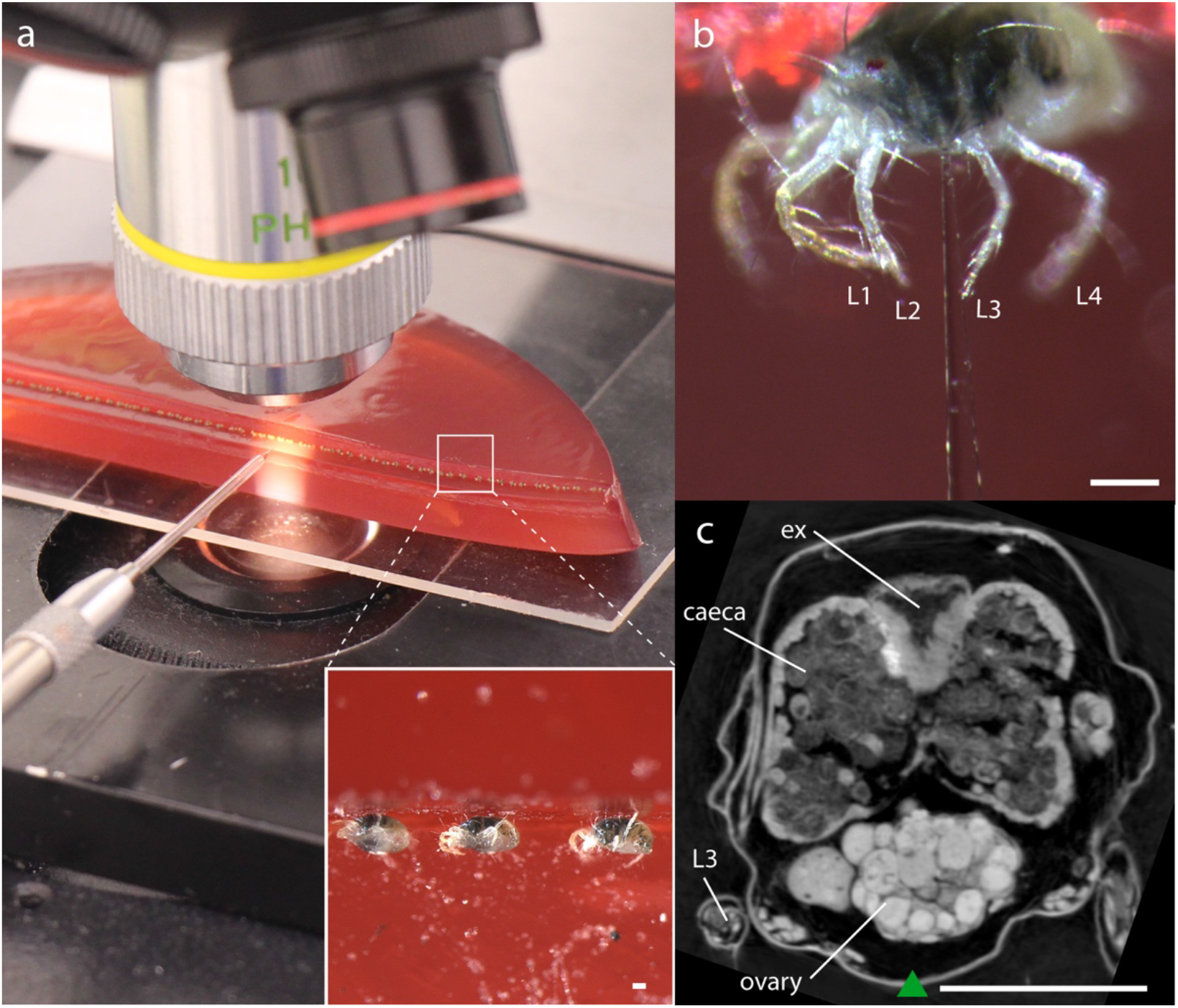
Cas9-sgRNA micro-injection setup for *T. urticae*. (a) setup for injection of *T. urticae* females: virgin females were aligned on an “agar platform” and injected under a microscope; insert: mites aligned on the agar platform, (b) females approximately injected at the third pair of legs: L1, L2, L3 and L4 refer to the 1^st^, 2^nd^, 3^rd^ and 4^th^ pair of legs, respectively (c) virtual cross-section at the third pair of legs; this section was obtained from a previously performed submicron CT scan of a *T. urticae* adult female (Jonckheere et al., 2016); a green triangle points towards Cas9-sgRNA injection location; L3: third pair of legs, ex= excretory organ. Scale bar in each panel represents 0.1 mm.

**Figure 2.**
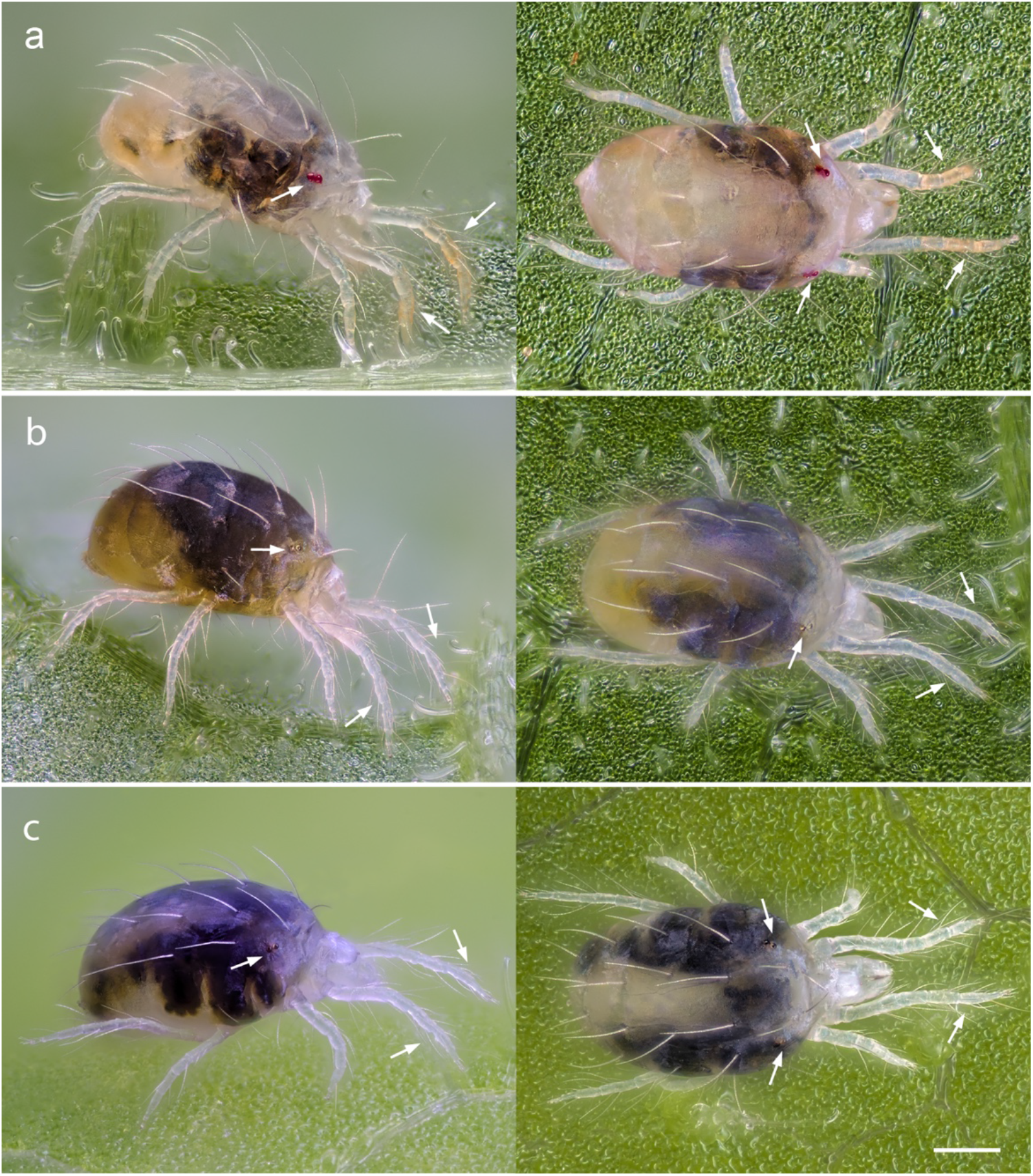
Phenotypes of adult *T. urticae* females of the WT strain, CRISPR line A and B. Shown are (a) *T. urticae* pigmentation of the WT strain, (b) albino phenotype of CRISPR line A and (c) albino phenotype of CRISPR line B. In all cases, adult females are shown. Arrows indicate red eye spots or distal red-orange pigmentation in the forelegs of WT mites, which are absent in albino females of line A, while females of line B have no red pigmentation in the forelegs but slight traces of red pigmentation (here barely visible) are present in the eyes. Left: lateral view; Right: dorsal view. Scale bar represents 0.1 mm.

### 2.5 Mode of inheritance of albino phenotype and generation of homozygous albino CRISPR lines A and B

Albino sons from RNP-injected females from the A and B batch were isolated on bean leaf disks (one male per leaf disk) and allowed to mate with three to five virgin females of the parental strain (London, WT). Mated females were allowed to lay eggs for six days on the leaf disk (disk 1) and were discarded afterwards. Next, three F_1_ teliochrysalis females that developed from eggs on disk 1, were transferred to a separate leaf disk, allowed to hatch, and to lay eggs for four days (disk 2). These virgin F_1_ females (from disk 2) were then transferred to another leaf disk and kept at 10°C to increase their life span (disk 3). Subsequently, the number of albino and WT males was counted on disk 2 and an albino male from disk 2 was mated with its virgin mother (on disk 3) to generate a homozygous albino line (CRISPR lines A and B). For these two lines, we also performed a complementation test on detached bean leaves. Briefly, 15 virgin (teliochrysalis) females from CRISPR line A or B were crossed with 30 males from the Alb-NL strain (Bryon et al., 2017). At least 100 resulting F_1_ females were assessed for albinism. Last, we also performed a complementation test between 15 teliochrysalis females of CRISPR line A and 30 males of CRISPR line B, and scored albinism for at least 100 F_1_ females.

### 2.6 DNA and RNA extraction from *T. urticae* CRISPR lines A and B and PCR amplification of *tetur01g11270*

DNA was collected from five pooled females from lines A and B using the CTAB method previously described by Navajas et al. (1998). PCR of *tetur01g11270* fragments was performed using the primers of the *in vitro* Cas9-sgRNA cleavage experiment (Table S1) and extracted DNA from lines A and B was used as template. The reactions consisted of 3 μl 10x Buffer, 0.2 mM of each dNTP, 0.33 μM of each primer, 2 μl template, 1U Kapa Taq DNA Polymerase (Kapa Biosystems) in a final volume of 30 μl and with cycling conditions as follows: 5 min at 95°C followed by 40 cycles of 30 s at 95°C, 40 s at 53°C, 1 min at 72°C and a final extension of 2 min at 72°C. PCR amplicons were verified on a 1.5% agarose gel, purified using the NucleoSpin^®^ Gel and PCR Clean-Up kit (Macherey-Nagel) according to the manufacturers’ instructions. Nucleotide sequences were determined in both strands of purified PCR products at the CeMIA sequencing facility (CEMIA, SA., Greece). Finally, RNA was extracted from mites of the A and B line. About 100 females were collected and RNA was extracted using the Qiagen RNeasy PLUS Kit (Qiagen Benelux, Venlo, Nederland). One μg of total RNA was used as template for synthesizing cDNA with the Maxima First Strand cDNA synthesis Kit for RT-qPCR (Fermentas Life Sciences, Aalst, Belgium). Primer3 (Rozen and Skaletsky, 2000) was used to design primers (tetur01g11270_cDNA primers) that amplify the coding sequence of the phytoene desaturase gene (*tetur01g11270*) (Table S1. PCRs were performed using the Expand Long Range dNTP Pack (Roche/Sigma-Aldrich, Belgium). Reaction mixtures were prepared according to the manufacturer’s instructions. The thermal profile consisted of denaturation for 2 min at 92°C, followed by 4 touch-down cycles of denaturation at 92°C for 10 s, annealing at 57°C −1°C/cycle for 15 s and elongation at 68°C for 2.5 min. Next, 40 cycles of 92°C for 10 s, 53°C for 15 s and 68°C for 2.5 min. After a final elongation of 68°C for 7 min, PCR products were purified using the E.Z.N.A. Cycle Pure kit (Omega Biotek) and Sanger sequenced by LGC genomics (Germany) with forward and reverse primers and four internal primers (Table S1).

### 2.7 Imaging

Images of adult females and immature stages of *T. urticae* were taken with an Olympus OM-D E-M1 mark II using a micro-objective on bellows (Nikon PB- 4). The following micro-objectives were used: a Nikon M Plan 10x 160/0.25 (for females and larvae of WT strain and CRISPR line A), Nikon achromatic 10x 160/0.25 (for females of CRISPR line B) and a Nikon BD Plan ELWD 20x 210/0.4 (for larvae of CRISPR line B). Between 50-150 pictures were used for a focus stack. The open-source software align_image_stack (https://www.systutorials.com/docs/linux/man/1-align_image_stack/) and Enfuse (http://software.bergmark.com/enfuseGUI/Main.html) were used to generate the focus stack, while Darktable (https://www.darktable.org/) was used for pre-and posttreatment of images. Images of adult males were taken using a stereomicroscope (Leica S8 Apo, Witzlar Germany) and a Leica DFC295 camera.

## 3 Results

### 3.1 sgRNA guide sequence design and *in vitro* Cas9-sgRNA cleavage

Guide sequences were designed using the CRISPOR website as described above. The first guide sequence (g1, 5’-GGTGGCAAGAGCACGAGCAC-3’) was selected because it had the highest “out-of-frame” score (the higher this score, the more deletions have a length that is not a multiple of three (Bae et al., 2014)) while the other guide sequence (g2, 5’-ACAATGGGTACTCCAGTACC-3’) was selected because it was located in a region postulated to encode the carotenoid binding domain of the phytoene desaturase (Armstrong et al., 1989; Sanz et al., 2002). Finally, both guide sequences had a predicted off-target count of zero. *In vitro* Cas9-sgRNA cleavage of PCR amplicons of *tetur01g11270* resulted in the correct *in silico*-predicted digestion pattern: amplicon 1 (895 bp) was cleaved into a 537 and 398 bp fragment, while amplicon 2 (699bp) was cleaved into a 197 bp and 502 bp fragment (Figure S3).

### 3.2 *In vivo* Cas9-sgRNA experiment

#### 3.2.1 Screening of albino male progeny and generation of CRISPR lines A and B

Two batches of virgin females were injected in the ovary: 245 mites in batch “A” and 177 mites in batch “B”. Twenty-four hours after injection, the percentage of alive females was recorded as 78.4% and 71.8%, respectively. Injected females were allowed to lay eggs for 24h, were placed on new arenas, and allowed to lay eggs for another 24 hours. The number of eggs on each arena was, approximately, 650 and 900 for batch A and 260 and 650 for batch B after 24 h and 24-48 h, respectively. After hatching, we screened for male larvae lacking pigment. In the arenas with eggs deposited within 24 hours after injection, we found one alive albino male in both batch A and B (Table 1), while in batch A thirteen specimens with albino phenotype were detected in larvae/protochrysalises resulting from eggs deposited between 24 and 48 hours after injection. However, none of these larvae/protochrysalises developed into adults. From both batches, the alive albino male was isolated, allowed to develop to the adult stage and crossed to obtain homozygous stable lines named CRISPR line A and B, respectively, which were characterized further. All life stages of CRISPR line A lacked red pigments (Figure 3, Figure S4). In contrast, only immature stages lacked red pigmentation in CRISPR line B, while adult stages do show traces of red pigmentation in the eyes, especially visible in the males, but lack red pigmentation in the forelegs (Figure 3, Figure S4, Figure S5).

**Table 1.** CRISPR-Cas9 efficiency.

|  | injection batch |  |
| --- | --- | --- |
|  | A | B |
| number of injected virgin females | 245 | 177 |
| number of injected virgin females alive after 24h | 192 | 127 |
| number of CRISPRed albino male offspring alive* | 1 | 1 |
| number of CRISPRed albino male offspring not alive** | 13 | 0 |
| % CRISPR-Cas9 success | 5.71 | 0.56 |

\*all alive offspring were found in the first 24h after injection
\*\*assuming albino phenotype is caused by a CRISPR-Cas9 event; all collected dead offspring were either in the protonymph or protochrysalis stage

**Figure 3.**
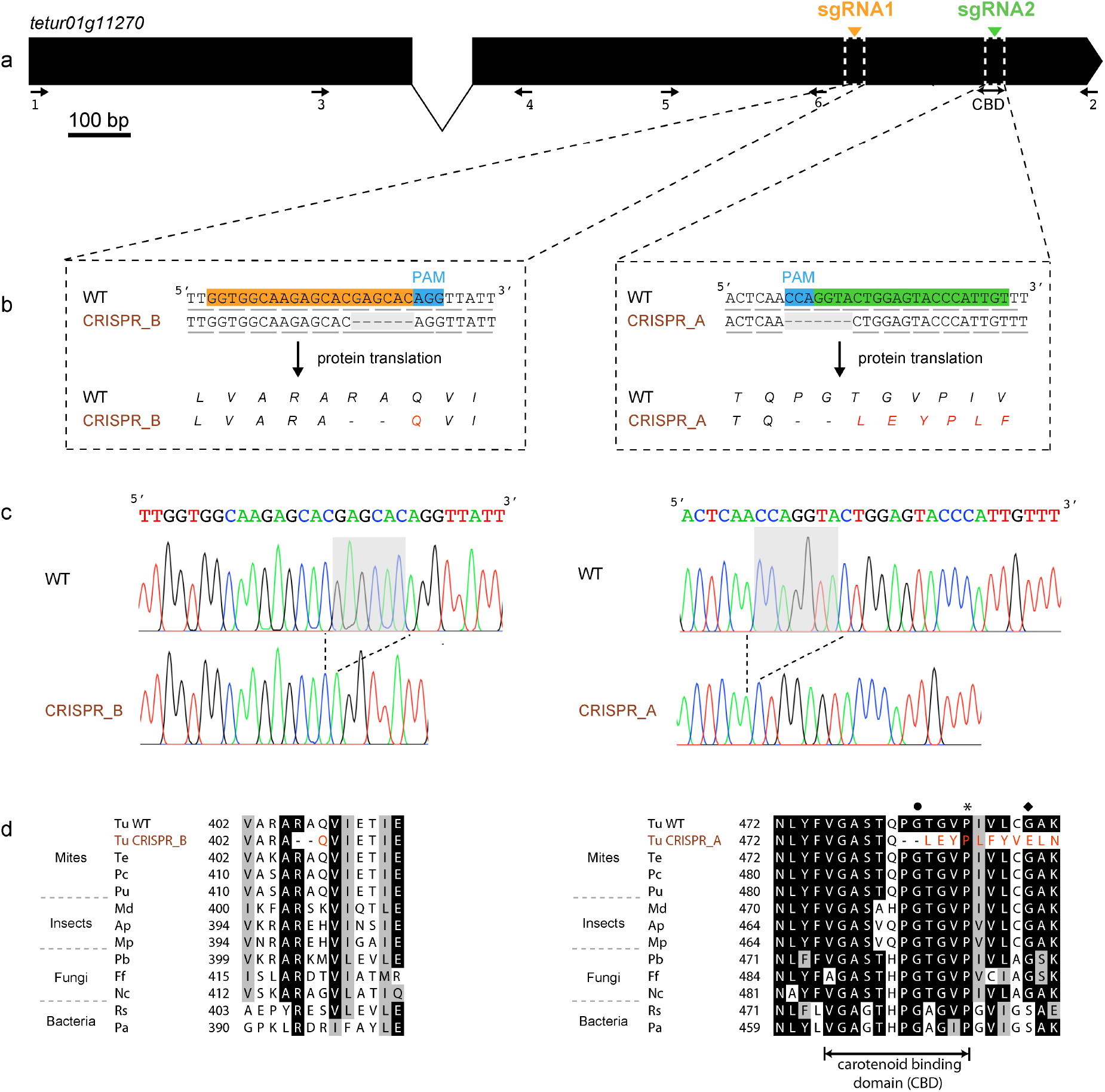
Small indels detected in the phytoene desaturase gene (*tetur01g11270*) of *T. urticae* females of CRISPR line A and B. (a) gene structure of *tetur01g11270*; the position of sgRNA1 and sgRNA2 cutting sites are indicated with an orange and green triangle, respectively; the position of the primers (1-6) used for PCR and sequencing of *tetur01g11270* cDNA is indicated with arrows (Table S2); (b) indels found adjacent to the sgRNA cutting sites in *tetur01g11270* of females of CRISPR line A or B; the guide sequence of sgRNA1 and the reverse complement of the sgRNA2 guide sequence are highlighted in orange and green, respectively, while the protospacer adjacent motif (PAM) is highlighted in blue; codons are underlined; (b-left) a 6 bp deletion (shaded gray) was found in *tetur01g11270* of females of CRISPR line B, resulting in the deletion of two amino acids; (b-right) a 7 bp deletion (shaded gray) was found in females from CRISPR line A, causing a deletion of two amino acids in the carotenoid binding domain and a frame shift changing translation; (c) chromatogram of the sequences displayed in (b), with the deletions present in the CRISPR lines shaded gray; (d) alignment of *tetur01g11270* of CRISPR line B (d-left) and CRISPR line A (d-right) with those of other tetranychid mites (*Te, Tetranychus evansi*, *Pu, Panonychus citri, Pc, Panonychus ulmi*), insects (*Md*, *Mayetiola destructor*, *Ap*, *Acyrthosiphon pisum, Mp*, *Myzus persicae*), Fungi *(Pb*, *Phycomyces blakesleeanus*, *Ff*, *Fusarium fujikuroi* and *Nc*, *Neurospora crassa*) and Bacteria (*Rs, Rhodobacter sphaeroides* and *Pa*, *Pantoea ananatis*). Accession numbers of all sequences can be found in Bryon et al. (2017) and in Supplementary Figure S6. (d-right) Mutations in *P. blakesleeanus* and *F. fujikiroi* that result in lowered phytoene desaturase activities are indicated with a black dot and rhombus, respectively (Prado-Cabrero et al., 2009; Sanz et al., 2002), while a Pro487Leu mutation that was identified in *tetur01g11270* of *T. urticae* lines W-Alb-1/W-Alb-2, with young stages lacking pigment but red color being apparent in adults (Bryon et al., 2017), is indicated with an asterisk.

#### 3.2.2 Mode of inheritance and complementation test of albino phenotype in CRISPR lines A and B

The genetic basis of the albino phenotype found in males of CRISPR lines A and B was determined by crossing line A and B males with females of the original WT strain. In all cases, F_1_ females of the resulting cross had normal body and eye color (Table 2). Together with the finding of an approximate 1:1 ratio of albino to WT phenotype in haploid F_2_ sons produced by virgin F1 females, this strongly indicated that albinism was inherited as a monogenic recessive trait. In a complementation test, females of CRISPR line A and males of CRISPR line B were crossed, and the resulting F_1_ females were all albinos indicating that the albino phenotype in both lines is caused by a disruption in the same gene (Table 2). Finally, we also crossed females of CRISPR lines A and B with males of strain Alb-NL, known to have an inactivating mutation in the phytoene desaturase gene (*tetur01g11270*) (Bryon et al. 2017), and found that all female F_1_ progeny was albino. This failure to complement suggests that the albino phenotype of CRISPR lines A and B results from a mutation or disruption in *tetur01g11270*, the gene targeted by our Cas9-sgRNA experiment.

**Table 2.** Inheritance and complementation tests.

| Crosses | F<sub>1</sub> ♂, % albino? | F<sub>2</sub> haploid males | | $\chi^2$ | P value |
| --- | --- | --- | --- | --- | --- |
|  |  | ALB | WT |  |  |
| *Inheritance tests (female x male)\** |  |  |  |  |  |
| WT x CRISPR A (rep1) | 0 | 15 | 20 | 0.714 | 0.39802 |
| WT x CRISPR A (rep2) | 0 | 30 | 21 | 1.588 | 0.20758 |
| WT x CRISPR A (rep3) | 0 | 17 | 20 | 0.243 | 0.62187 |
| WT x CRISPR B (rep1) | 0 | 25 | 20 | 0.556 | 0.45606 |
| WT x CRISPR B (rep2) | 0 | 12 | 14 | 0.154 | 0.69489 |
| WT x CRISPR B (rep3) | 0 | 28 | 27 | 0.018 | 0.89274 |
| *Complementation tests (female x male)\*\** |  |  |  |  |  |
| CRISPR A x Alb-NL*** | 100 |  |  |  |  |
| CRISPR B x Alb-NL | 100 |  |  |  |  |
| CRISPR A x CRISPR B | 100 |  |  |  |  |

\*an alive albino male - CRISPR A or CRISPR B - that was detected in the progeny of Cas9-sgRNA injected females of either batch A or B, respectively, was crossed with three to five females of the WT strain (1 male x 3-5 females) in 3 replicates
\*\* 15 females crossed with 30 males; 100 F<sub>1</sub> females were screened for wildtype or albino phenotype
\*\*\* the Alb-NL strain is an albino *T. urticae* strain known to have an inactivating mutation in its carotenoid desaturase gene (*tetur01g11270*) (Bryon et al. 2017)

#### 3.2.3 Sequence analysis of *tetu01g11270* in CRISPR lines A and B

DNA was extracted from CRISPR lines A and B and sequencing of PCR amplicons 1 and 2 revealed disruptions in the *tetur01g11270* gene in both lines. *Tetur01g11270* of CRISRP line B harbored a 6 bp deletion (nt 1117-1122 in WT reference sequence of *tetur01g11270*) that was located 6 bp upstream of the sgRNA1 PAM site, causing a loss of two amino acids (Arg406 and Ala407).

Based on an alignment of phytoene desaturases of insects, fungi and bacteria (Figure 3d) Arg406 is highly conserved. CRISPR line A harbored a 7 bp deletion (nt 1444-1450 in *tetur01g11270* in WT reference sequence of *tetur01g11270*) that was located 4 bp upstream of the sgRNA2 PAM site, resulting in the loss of two amino acids and a frame shift, changing translation (Figure 3b) in the region of the carotenoid binding domain (Armstrong et al., 1989). To assure that the detected deletions were the only disruptions in the coding sequence of *tetur01g11270* of CRISPR lines A and B, we sequenced the complete cDNA sequence of *tetur01g11270* of both CRISPR lines and the WT strain. The cDNA sequence of CRISPR line B was, except for the 6 bp deletion, 100% identical to that of the WT strain, while in the cDNA sequence of CRISPR line A, we found, next to the 7 bp deletion, three non-synonymous single nucleotide polymorphisms (SNPs) (Figure S6). All three non-synonymous SNPs resulted in favored substitutions according to Russel et al. (2003). The amino acid changes “K->Q” and “I->V” (Figure S6), caused by two non-synonymous SNPs, occur at a non-conserved amino acid position in the phytoene desaturase protein (Figure S6 and Supplemental Figure S5 in Bryon et al. Bryon et al. (2017)) and were also present in the WT strain at low frequency (data not shown). Last, the remaining non-synonymous SNP (resulting in an amino acid change “V->I”) was located downstream of the 7 bp deletion.

## 4 Discussion

CRISPR-Cas9 has revolutionized genome editing in metazoan species, including more and more arthropods (Kotwica-Rolinska et al., 2019; Reardon, 2019; Sun et al., 2017). For many arthropods, the ortholog of the *Drosophila white* gene, an ABC-transporter essential for eye pigmentation, has been used as a CRISPR-Cas9 target for establishing proof of principle of this technology (Bai et al., 2019; Ismail et al., 2018; Khan et al., 2017; Xue et al., 2018). For chelicerates, such as mites and ticks, however, the CRISPR-Cas9 technology has not yet been validated. The main reason is probably because the injection of mite and tick embryos is currently not feasible (Khila and Grbić, 2007; Sharma, 2017) and because non-lethal convenient genetic markers with a clearly visible phenotype are not yet available. In the two-spotted spider mite *T. urticae*, for example, a clear ortholog of the *white* gene could not be identified (Dermauw et al. 2013). However, recent studies have uncovered several mutations that result in pigmentation defects in a number of spider mite populations and species (Bryon et al., 2017; Wybouw et al., 2019b). For example, in *T. urticae*, it was shown that several mutations in a gene encoding a phytoene desaturase (*tetur01g11270*) caused an albino phenotype (lack of red pigment in frontal legs and eyes) (Bryon et al., 2017). Interestingly, like in a few other arthropods (Zhao and Nabity, 2017), this gene was horizontally acquired from fungi (Altincicek et al., 2012; Grbic et al., 2011) and encodes an enzyme that catalyzes the formation of lycopene, from which β-carotene and other red pigments are derived (Maoka, 2019). The discovery of a *T. urticae* genetic marker with a clearly visible phenotype, scorable in larvae and even embryos, significantly facilitates screening for potential genetic transformants. We therefore took advantage of this discovery to design a CRISPR-Cas9 strategy with sgRNAs that target the phytoene desaturase of *T. urticae* (Figure 3). Next to the availability of a genetic marker with a clearly visible phenotype, efficient CRISPR-Cas9 further requires the delivery of the Cas9-sgRNA complex into the embryos in early development. As successful injection of mite and tick embryos has currently not been achieved (see above), we followed a strategy previously applied for nematodes, mosquitoes and psyllids (Chaverra-Rodriguez et al., 2018; Cho et al., 2013; Hunter et al., 2018; Macias et al., 2019), and we injected *T. urticae* females in the ovary, assuming that the Cas9-sgRNA complex would be incorporated into the oocytes and developing embryos. In addition, the arrhenotokous reproduction system allowed us to inject unfertilized females of which the progeny consists of haploid males only. This allowed to immediately screen for an albino phenotype among the male progeny of injected females.

In this study, two batches (A and B) of virgin *T. urticae* females were injected with Cas9-sgRNA and in each batch one albino male was identified in the progeny developed from eggs laid by females less than 24 hours after injection (Table 1). Subsequently, homozygous albino lines (CRISPR line A and B) were generated from these males and both the mode of inheritance and the complementation test revealed that disruptions in *tetur01g11270* caused the albino phenotype (Table 2). The *T. urticae* genome harbors three copies of phytoene desaturase, and although *tetur01g11270* is the only one with a clear role in pigment synthesis (Bryon et al., 2013), one could question whether other *T. urticae* phytoene desaturase genes (*tetur11g04820* and *tetur11g04810*, Grbić et al. (2011)) were also targeted. However, complementation tests with a characterized albino line point to a single causal gene (Table 2). In addition, no off-target effects were predicted for guide sequences of both sgRNAs, and guide sequence regions differ significantly between *tetur01g11270* and the other two phytoene desaturase genes (Figure S7). Further, to assess whether the *tetur01g11270* disruptions were caused by typical CRISPR-Cas9 events, we sequenced *tetur01g11270* of CRISPR lines A and B at the DNA and cDNA level. Typical CRISPR-Cas9 events (Jinek et al., 2012) were identified in *tetur01g11270* of both lines, with deletions located four to six base pairs upstream of the PAM site (Figure 3). Sequencing of the *tetur01g11270* full-length coding sequence revealed that no other polymorphisms could be detected in CRISPR line B compared to the WT strain, while the *tetur01g11270* coding sequence of CRISPR line A did contain three favored non-synonymous mutations (Figure S6) of which two were also present in the WT strain. Altogether, this leaves no doubt that the Cas9-induced deletions in *tetur01g11270* of CRISPR lines A and B are the underlying genetic basis of the albino phenotype. Subtle differences in the albino phenotype of each line could to some extent also be linked to the type of the Cas9-sgRNA induced deletion. In CRISPR line A, the 7 bp deletion in *tetur01g11270* causes a frameshift, thereby abolishing the carotenoid binding domain (Armstrong et al., 1989), resulting in the lack of pigment in all stages. In CRISPR line B, the 6 bp deletion results in the loss of two amino acids, including a highly conserved arginine, but does not change translation (Figure 3d). While immature stages of CRISPR line B lack pigmentation, the eyes of adult females and especially males of CRISPR line B, traces of red pigmentation could be observed, suggesting the 6 bp deletion can be considered as a hypomorphic mutation, i.e. causing only a partial loss of gene function (Muller, 1932).

Based on the total number of eggs that was laid by the injected females (1550 and 910 for batches A and B, respectively), the percentage of CRISPR-Cas9 transformed mite embryos is low (Table 1). Especially when compared to the CRISPR-Cas9 efficiency in nematodes, where a mutation frequency of up to 17% in the F_1_ progeny can be obtained by injection of the Cas9-sgRNA complex into the gonads (Cho et al., 2013). Furthermore, in contrast to the 24h egg arenas of batch A and B, we could not obtain alive albino males from the 24-48h egg arena of batch A, as all thirteen detected albino larvae/protochrysalises did not develop into adulthood. Although genetic evidence was not gathered, we can assume that the observed albino males with identical phenotype in interval 24-48h were caused by CRISPR-Cas9 events. If this is the case, the decreased survival of the larvae/protochrysalises might have been the result of multiple accompanying off-target CRISPR-Cas9 events at this time point after injection. However, given that the CRISPOR software predicted that both sgRNAs have zero off-target effects, this seems unlikely. If only the 24h time point is taken into consideration, we obtained about one transformant per 200 injected females, a frequency that does allow to screen for visible phenotypic traits immediately. Arrhenotokous reproduction allows to immediately screen the males that can be directly used in dedicated crosses to fix the mutation. Which time point after injection is the most likely to result in CRISPRed embryos should be investigated and optimization of this timing could potentially increase screening efficacy. Because of this straightforward phenotype screening and mutation fixation, a ‘CO-CRISPR’ approach might be used to make this strategy also feasible for mutations without a visible phenotype. In this approach, injection mixtures would contain sgRNA for both a marker gene and additional target-gene. It was previously shown for nematodes that transformants with the visible marker have a much higher frequency of mutations in the target-gene (Dickinson and Goldstein, 2016; Farboud et al., 2019). This allows to preselect a number of progeny for further screening.

Previously, Bryon et al. (2017) used a similar CRISPR-Cas9 approach in an attempt to provide functional evidence of the role of mutations and deletions in *tetur01g11270* in albinism. However, typical CRISPR-Cas9 events were not recorded. We hypothesized that this was most likely due to insufficient RNP uptake by the oocytes. Here, we increased the Cas9 protein concentration more than 5-fold to 4.85 μg/μl. Furthermore, we also added chloroquine, because it was recently shown that the addition of this compound improves CRISPR-Cas9 efficiency in mosquitoes (Chaverra-Rodriguez et al. 2018). Recent studies also hint toward other modifications that could improve CRISPR-Cas9 transformation efficiency, such as the use of other adjuvants like lipofectamine or branched amphiphilic peptide capsules (BAPC) (Adams et al., 2019; Hunter et al., 2018), or a shorter Cas protein (Rusk, 2019). Last, in a recent breakthrough study it was shown how ReMOT (Receptor-Mediated Ovary Transduction of Cargo) can be exploited to deliver Cas9 in oocytes after the injection of female mosquitoes. In this system, a “guide peptide” (P2C) mediates the transduction of the Cas9 RNP complex from the female mosquito hemolymph to developing oocytes. Although the principle of transformation should be transferable to other organisms, the peptide and protein identified in Chaverra-Rodriguez et al. (2018) have no homologs outside dipterans (flies and mosquitoes) and might not be readily transferable to mites and ticks.

In conclusion, two independent mutagenesis events were induced in the spider mite *T. urticae* using CRISPR-Cas9, providing an impetus for genetic transformation in chelicerates and paving the way for functional studies using CRISPR-Cas9 in *T. urticae*.

## Author contributions

WD and TVL designed experiments; WD, WJ, MR and IL performed experiments. WD and TVL wrote the manuscript, with input from JV, MR and WJ. All authors reviewed the manuscript.

## Acknowledgements

We thank Merijn Kant (University of Amsterdam, The Netherlands) for providing the Alb-NL strain, Gilles San Martin (Walloon Agricultural Research Centre CRA-W, Gembloux, Belgium) for taking photographs (Figure 2, Figure S4) of the different spider mite lines, Astrid Bryon (University of Wageningen, The Netherlands) for providing Figure 1a and René Feyereisen (University of Copenhagen, Denmark/University of Ghent, Belgium) for critical reading of the manuscript. This work was supported by the European Union’s Horizon 2020 research and innovation program [grant 772026-POLYADAPT to TVL and 773902-SuperPests to TVL and JV]. During this study WD was a postdoctoral fellow of the Research Foundation Flanders (FWO).

## Supplementary Figure Legends

**Figure S1 - Agar platforms used for injection of *T. urticae* females**

(a) two microscopic slides attached to each other; (b) cherry/agar plate containing the two microscope slides; after solidification of agar, slides were removed from the agar and the agar plate was cut in two along the length of the slide indentation.

**Figure S2 - Injection needle used for injections of *T. urticae* females.** (a) injection needle pulled from Clark capillary glass; scale bar represents 0.1 mm (b) close-up of the tip of the pulled needle.

**Figure S3 - *In vitro* digestion with Cas9/sgRNAs of two PCR amplicons of *tetur01g11270* of adult females of the *T. urticae* WT strain.** lane 1: Cas9 with PCR amplicon 1 (895 bp); lane 2: Cas9 with PCR amplicon 1 <u>and</u> sgRNA1, resulting in a 537 bp and 398 bp fragment (black arrows); lane 3: Cas9 with PCR amplicon 2 (699 bp); lane 4: Cas9 with PCR amplicon 2 <u>and</u> sgRNA2 resulting in a 502 bp and 197 bp fragment (white arrows); M: BenchTop 100 bp DNA ladder (Promega, catalog# G8291).

**Figure S4 - Phenotype of immature stages of the *T. urticae* WT strain and CRISPR lines A and B.** Shown are (a) *T. urticae* pigmentation of the WT strain, (b) albino phenotype of CRISPR line A and (c) albino phenotype of CRISPR line B. In (a) and (c) larval stages are shown while in (b) a protonymphal stage is shown. Arrows indicate red eye spots of WT mites that are absent in immature stages of line A and B. Scale bar represents 0.1 mm.

**Figure S5 - Phenotype of adult males of the *T. urticae* WT strain and CRISPR lines A and B**. Shown are (a) *T. urticae* pigmentation of the WT strain, (b) albino phenotype of CRISPR line A and (c) albino phenotype of CRISPR line B. Arrows indicate red eye spots of WT mites that are absent in males of line A, while traces of red pigment can be seen in the eyes of males of line B. Scale bar represents 0.1 mm.

**Figure S6 - Nucleotide alignment of cDNA of *tetur01g11270* of the *T. urticae* WT strain and CRISPR line A and B**. Nucleotides with 100% identity are shaded black; *tetur01g11270* of CRISPR B line was completely identical to *tetur01g11270* of the WT strain while three non-synonymous SNPs (indicated in blue font) were found in *tetur01g11270* cDNA of the A strain.

**Figure S7 - Alignment of *tetur01g11270*, *tetur11g04810* and *tetur11g04820* of the London strain (Grbic et al. 2011) with guide sequences of sgRNA1 and sgRNA2**. Guide sequences of sgRNA1 and sgRNA2 are shaded orange and green respectively.

## Supplementary Tables

**Table S1 - Composition of CRISPR-Cas9 injection mix**

**Table S2 - Primers used in this study**

## References

Adams, S., Pathak, P., Shao, H., Lok, J.B., Pires-daSilva, A., 2019. Liposome-based transfection enhances RNAi and CRISPR-mediated mutagenesis in non-model nematode systems. Sci Rep 9, 483.

Alba, J.M., Schimmel, B.C.J., Glas, J.J., Ataide, L.M.S., Pappas, M.L., Villarroel, C.A., Schuurink, R.C., Sabelis, M.W., Kant, M.R., 2015. Spider mites suppress tomato defenses downstream of jasmonate and salicylate independently of hormonal crosstalk. New Phytol 205, 828–840.

Altincicek, B., Kovacs, J.L., Gerardo, N.M., 2012. Horizontally transferred fungal carotenoid genes in the two-spotted spider mite *Tetranychus urticae*. Biology Letters 8, 253–257.

Armstrong, G.A., Alberti, M., Leach, F., Hearst, J.E., 1989. Nucleotide sequence, organization, and nature of the protein products of the carotenoid biosynthesis gene cluster of *Rhodobacter capsulatus*. Molecular and General Genetics MGG 216, 254–268.

Bae, S., Kweon, J., Kim, H.S., Kim, J.-S., 2014. Microhomology-based choice of Cas9 nuclease target sites. Nat Meth 11, 705–706.

Bai, X., Zeng, T., Ni, X.-Y., Su, H.-A., Huang, J., Ye, G.-Y., Lu, Y.-Y., Qi, Y.-X., 2019. CRISPR/Cas9-mediated knockout of the eye pigmentation gene white leads to alterations in colour of head spots in the oriental fruit fly, *Bactrocera dorsalis*. Insect Mol Biol 28, 837–849.

Bajda, S., Dermauw, W., Panteleri, R., Sugimoto, N., Douris, V., Tirry, L., Osakabe, M., Vontas, J., Van Leeuwen, T., 2017. A mutation in the PSST homologue of complex I (NADH:ubiquinone oxidoreductase) from *Tetranychus urticae* is associated with resistance to METI acaricides. Insect Biochem Mol Biol 80, 79–90.

Betts, M.J., Russell, R.B., 2003. Amino acid properties and consequences of subsitutions in: Barnes, M.R., Gray, I.C. (Eds.), Bioinformatics for Geneticists,. Wiley.

Blaazer, C.J.H., Villacis-Perez, E.A., Chafi, R., Van Leeuwen, T., Kant, M.R., Schimmel, B.C.J., 2018. Why Do Herbivorous Mites Suppress Plant Defenses? Front Plant Sci 9, 1057.

Bryon, A., Kurlovs, A.H., Dermauw, W., Greenhalgh, R., Riga, M., Grbić, M., Tirry, L., Osakabe, M., Vontas, J., Clark, R.M., Van Leeuwen, T., 2017. Disruption of a horizontally transferred phytoene desaturase abolishes carotenoid accumulation and diapause in *Tetranychus urticae*. Proc Natl Acad Sci U S A 114, E5871–E5880.

Bryon, A., Wybouw, N., Dermauw, W., Tirry, L., Van Leeuwen, T., 2013. Genome wide gene-expression analysis of facultative reproductive diapause in the two-spotted spider mite *Tetranychus urticae*. BMC Genomics 14, 815.

Bui, H., Greenhalgh, R., Ruckert, A., Gill, G.S., Lee, S., Ramirez, R.A., Clark, R.M., 2018. Generalist and Specialist Mite Herbivores Induce Similar Defense Responses in Maize and Barley but Differ in Susceptibility to Benzoxazinoids. Front Plant Sci 9.

Chaverra-Rodriguez, D., Macias, V.M., Hughes, G.L., Pujhari, S., Suzuki, Y., Peterson, D.R., Kim, D., McKeand, S., Rasgon, J.L., 2018. Targeted delivery of CRISPR-Cas9 ribonucleoprotein into arthropod ovaries for heritable germline gene editing. Nat Comm 9, 3008.

Cho, S.W., Lee, J., Carroll, D., Kim, J.-S., Lee, J., 2013. Heritable Gene Knockout in *Caenorhabditis elegans* by Direct Injection of Cas9–sgRNA Ribonucleoproteins. Genetics 195, 1177–1180.

Concordet, J.-P., Haeussler, M., 2018. CRISPOR: intuitive guide selection for CRISPR/Cas9 genome editing experiments and screens. Nucleic Acids Res 46, W242–W245.

Dermauw, W., Osborne, E.J., Clark, R.M., Grbic, M., Tirry, L., Van Leeuwen, T., 2013a. A burst of ABC genes in the genome of the polyphagous spider mite *Tetranychus urticae*. BMC Genomics 14, 317.

Dermauw, W., Wybouw, N., Rombauts, S., Menten, B., Vontas, J., Grbic, M., Clark, R.M., Feyereisen, R., Van Leeuwen, T., 2013b. A link between host plant adaptation and pesticide resistance in the polyphagous spider mite *Tetranychus urticae*. Proc Natl Acad Sci U S A 110, E113–E122.

Dickinson, D.J., Goldstein, B., 2016. CRISPR-Based Methods for *Caenorhabditis elegans* Genome Engineering. Genetics 202, 885–901.

Douris, V., Steinbach, D., Panteleri, R., Livadaras, I., Pickett, J.A., Van Leeuwen, T., Nauen, R., Vontas, J., 2016. Resistance mutation conserved between insects and mites unravels the benzoylurea insecticide mode of action on chitin biosynthesis. Proc Natl Acad Sci U S A 113, 14692–14697.

Farboud, B., Severson, A.F., Meyer, B.J., 2019. Strategies for Efficient Genome Editing Using CRISPR-Cas9. Genetics 211, 431–457.

Garb, J.E., Sharma, P.P., Ayoub, N.A., 2018. Recent progress and prospects for advancing arachnid genomics. Curr Opin Insect Sci 25, 51–57.

Grbic, M., Van Leeuwen, T., Clark, R.M., Rombauts, S., Rouze, P., Grbic, V., Osborne, E.J., Dermauw, W., Ngoc, P.C.T., Ortego, F., Hernandez-Crespo, P., Diaz, I., Martinez, M., Navajas, M., Sucena, E., Magalhaes, S., Nagy, L., Pace, R.M., Djuranovic, S., Smagghe, G., Iga, M., Christiaens, O., Veenstra, J.A., Ewer, J., Mancilla Villalobos, R., Hutter, J.L., Hudson, S.D., Velez, M., Yi, S.V., Zeng, J., Pires-daSilva, A., Roch, F., Cazaux, M., Navarro, M., Zhurov, V., Acevedo, G., Bjelica, A., Fawcett, J.A., Bonnet, E., Martens, C., Baele, G., Wissler, L., Sanchez-Rodriguez, A., Tirry, L., Blais, C., Demeestere, K., Henz, S.R., Gregory, T.R., Mathieu, J., Verdon, L., Farinelli, L., Schmutz, J., Lindquist, E., Feyereisen, R., Van de Peer, Y., 2011. The genome of Tetranychus urticae reveals herbivorous pest adaptations. Nature 479, 487–492.

Gui, T., Zhang, J., Song, F., Sun, Y., Xie, S., Yu, K., Xiang, J., 2016. CRISPR/Cas9-Mediated Genome Editing and Mutagenesis of *EcChi4* in *Exopalaemon carinicauda*. G3 6, 3757–3764.

Hunter, W.B., Gonzalez, M.T., Tomich, J., 2018. BAPC-assisted CRISPR/Cas9 System: Targeted Delivery into Adult Ovaries for Heritable Germline Gene Editing (Arthropoda: Hemiptera). bioRxiv, 478743.

Iida, J., Desaki, Y., Hata, K., Uemura, T., Yasuno, A., Islam, M., Maffei, M.E., Ozawa, R., Nakajima, T., Galis, I., Arimura, G.-i., 2019. Tetranins: new putative spider mite elicitors of host plant defense. New Phytol 224, 875–885.

Ismail, N.I.B., Kato, Y., Matsuura, T., Watanabe, H., 2018. Generation of white-eyed *Daphnia magn*a mutants lacking scarlet function. PLoS One 13, e0205609.

Jinek, M., Chylinski, K., Fonfara, I., Hauer, M., Doudna, J.A., Charpentier, E., 2012. A Programmable Dual-RNA–Guided DNA Endonuclease in Adaptive Bacterial Immunity. Science 337, 816–821.

Jonckheere, W., Dermauw, W., Zhurov, V., Wybouw, N., Van den Bulcke, J., Villarroel, C.A., Greenhalgh, R., Grbić, M., Schuurink, R.C., Tirry, L., Baggerman, G., Clark, R.M., Kant, M.R., Vanholme, B., Menschaert, G., Van Leeuwen, T., 2016. The salivary protein repertoire of the polyphagous spider mite *Tetranychus urticae*: a quest for effectors. Mol Cell Proteomics 15, 3594–3613.

Khan, S.A., Reichelt, M., Heckel, D.G., 2017. Functional analysis of the ABCs of eye color in *Helicoverpa armigera* with CRISPR/Cas9-induced mutations. Sci Rep 7, 40025.

Khila, A., Grbić, M., 2007. Gene silencing in the spider mite *Tetranychus urticae*: dsRNA and siRNA parental silencing of the Distal-less gene. Dev Genes Evol 217, 241–251.

Korona, D., Koestler, S.A., Russell, S., 2017. Engineering the Drosophila Genome for Developmental Biology. Journal of developmental biology 5, 16.

Kotwica-Rolinska, J., Chodakova, L., Chvalova, D., Kristofova, L., Fenclova, I., Provaznik, J., Bertolutti, M., Wu, B.C.-H., Dolezel, D., 2019. CRISPR/Cas9 Genome Editing Introduction and Optimization in the Non-model Insect *Pyrrhocoris apterus*. Front Physiol 10.

Kurlovs, A.H., Snoeck, S., Kosterlitz, O., Van Leeuwen, T., Clark, R.M., 2019. Trait mapping in diverse arthropods by bulked segregant analysis. Curr Opin Insect Sci 36, 57–65.

Kwon, D.H., Park, J.H., Ashok, P.A., Lee, U., Lee, S.H., 2016. Screening of target genes for RNAi in *Tetranychus urticae* and RNAi toxicity enhancement by chimeric genes. Pestic Biochem Physiol 130, 1–7.

Macias, V.M., McKeand, S., Chaverra-Rodriguez, D., Hughes, G.L., Fazekas, A., Pujhari, S., Jasinskiene, N., James, A.A., Rasgon, J.L., 2019. Cas9-mediated gene-editing in the malaria mosquito *Anopheles stephensi* by ReMOT Control. bioRxiv, 775312.

Maoka, T., 2019. Carotenoids as natural functional pigments. Journal of Natural Medicines.

Martel, C., Zhurov, V., Navarro, M., Martinez, M., Cazaux, M., Auger, P., Migeon, A., Santamaria, M.E., Wybouw, N., Diaz, I., 2015. Tomato Whole Genome Transcriptional Response to *Tetranychus urticae* Identifies Divergence of Spider Mite-Induced Responses Between Tomato and Arabidopsis. Mol Plant Microbe Interact 28, 343–361.

Martin, A., Serano, J.M., Jarvis, E., Bruce, H.S., Wang, J., Ray, S., Barker, C.A., O’Connell, L.C., Patel, N.H., 2016. CRISPR/Cas9 mutagenesis reveals versatile roles of Hox genes in crustacean limb specification and evolution. Curr Biol 26, 14–26.

Mota-Sanchez, R.M., Wise, J.C., 2019. Arthropod Pesticide Resistance Database (APRD). Available at: https://www.pesticideresistance.org/.

Muller, H.J., 1932. Further studies on the nature and causes of gene mutations. Proceedings of the Sixth International Congress of Genetics, Ithaca, New York. 1, 213–255.

Nakanishi, T., Kato, Y., Matsuura, T., Watanabe, H., 2014. CRISPR/Cas-Mediated Targeted Mutagenesis in Daphnia magna. PLoS One 9, e98363.

Navajas, M., Lagnel, J., Gutierrez, J., Boursot, P., 1998. Species-wide homogeneity of nuclear ribosomal ITS2 sequences in the spider mite *Tetranychus urticae* contrasts with extensive mitochondrial COI polymorphism. Heredity 80, 742–752.

Prado-Cabrero, A., Schaub, P., Díaz-Sánchez, V., Estrada, A.F., Al-Babili, S., Avalos, J., 2009. Deviation of the neurosporaxanthin pathway towards β-carotene biosynthesis in *Fusarium fujikuroi* by a point mutation in the phytoene desaturase gene. The FEBS Journal 276, 4582–4597.

Presnail, J.K., Hoy, M.A., 1992. Stable genetic transformation of a beneficial arthropod, Metaseiulus occidentalis (Acari: Phytoseiidae), by a microinjection technique. Proc Natl Acad Sci U S A 89, 7732–7736.

Reardon, S., 2019. CRISPR gene-editing creates wave of exotic model organisms. Nature 568, 441–442

Riga, M., Bajda, S., Themistokleous, C., Papadaki, S., Palzewicz, M., Dermauw, W., Vontas, J., Leeuwen, T.V., 2017. The relative contribution of target-site mutations in complex acaricide resistant phenotypes as assessed by marker assisted backcrossing in *Tetranychus urticae*. Sci Rep 7, 9202.

Rozen, S., Skaletsky, H.J., 2000. Primer3 on the WWW for general users and for biologist programmers, in: Krawetz, S., Misener, S. (Eds.), Bioinformatics Methods and Protocols: Methods in Molecular Biology. Humana Press, Totowa, New Jersey, USA, pp. 365–386.

Rungger, D., Muster, L., Georgiev, O., Rungger-Brändle, E., 2017. Oocyte shuttle, a recombinant protein transporting donor DNA into the *Xenopus* oocyte *in situ*. Biology Open 6, 290–295.

Rusk, N., 2019. Spotlight on Cas12. Nat Meth 16, 215–215.

Santamaría, M.E., Hernández-Crespo, P., Ortego, F., Grbic, V., Grbic, M., Diaz, I., Martinez, M., 2012. Cysteine peptidases and their inhibitors in *Tetranychus urticae*: a comparative genomic approach. BMC Genomics 13, 307.

Santamaría, M.E., Martínez, M., Arnaiz, A., Rioja, C., Burow, M., Grbic, V., Díaz, I., 2019. An Arabidopsis TIR-Lectin Two-Domain Protein Confers Defense Properties against *Tetranychus urticae*. Plant Physiol 179, 1298–1314.

Sanz, C., Alvarez, M.I., Orejas, M., Velayos, A., Eslava, A.P., Benito, E.P., 2002. Interallelic complementation provides genetic evidence for the multimeric organization of the *Phycomyces blakesleeanus* phytoene dehydrogenase. Eur J Biochem 269, 902–908.

Schlachter, C.R., Daneshian, L., Amaya, J., Klapper, V., Wybouw, N., Borowski, T., Van Leeuwen, T., Grbic, V., Grbic, M., Makris, T.M., Chruszcz, M., 2019. Structural and functional characterization of an intradiol ring-cleavage dioxygenase from the polyphagous spider mite herbivore *Tetranychus urticae* Koch. Insect Biochem Mol Biol 107, doi:10.1016/j.ibmb.2018.1012.1001.

Scott, J.G., Michel, K., Bartholomay, L.C., Siegfried, B.D., Hunter, W.B., Smagghe, G., Zhu, K.Y., Douglas, A.E., 2013. Towards the elements of successful insect RNAi. J Insect Physiol 59, 1212–1221.

Scott, M.J., Gould, F., Lorenzen, M., Grubbs, N., Edwards, O., O’Brochta, D., 2018. Agricultural production: assessment of the potential use of Cas9-mediated gene drive systems for agricultural pest control. Journal of Responsible Innovation 5, S98–S120.

Sharma, A., 2017. Development of CRISPR-Cas9 gene drive system for deer tick, *Ixodes scapularis*, IGTRCN Peer-to-Peer Training Fellowship Report. Available at: http://igtrcn.org/wp-content/uploads/2018/01/Sharma_IGTRCN_report_val.docx, University of Maryland, MD, USA.

Snoeck, S., Wybouw, N., Van Leeuwen, T., Dermauw, W., 2018. Transcriptomic Plasticity in the Arthropod Generalist *Tetranychus urticae* Upon Long-Term Acclimation to Different Host Plants. G3 8, 3865–3879.

Sun, D., Guo, Z., Liu, Y., Zhang, Y., 2017. Progress and prospects of CRISPR/Cas systems in insects and other arthropods. Front Physiol 8, 608.

Suzuki, T., Nunes, M.A., España, M.U., Namin, H.H., Jin, P., Bensoussan, N., Zhurov, V., Rahman, T., De Clercq, R., Hilson, P., Grbic, V., Grbic, M., 2017. RNAi-based reverse genetics in the chelicerate model *Tetranychus urticae*: A comparative analysis of five methods for gene silencing. PLoS One 12, e0180654.

Van Leeuwen, T., Demaeght, P., Osborne, E.J., Dermauw, W., Gohlke, S., Nauen, R., Grbic, M., Tirry, L., Merzendorfer, H., Clark, R.M., 2012. Population bulk segregant mapping uncovers resistance mutations and the mode of action of a chitin synthesis inhibitor in arthropods. Proc Natl Acad Sci U S A 109, 4407–4412.

Van Leeuwen, T., Dermauw, W., 2016. The molecular evolution of xenobiotic metabolism and resistance in Chelicerate mites. Annu Rev Entomol 61, 475–498.

Van Leeuwen, T., Tirry, L., Yamamoto, A., Nauen, R., Dermauw, W., 2015. The economic importance of acaricides in the control of phytophagous mites and an update on recent acaricide mode of action research. Pestic Biochem Physiol 121, 12–21.

Villarroel, C.A., Jonckheere, W., Alba, J.M., Glas, J.J., Dermauw, W., Haring, M.A., Van Leeuwen, T., Schuurink, R.C., Kant, M.R., 2016. Salivary proteins of spider mites suppress defenses in *Nicotiana benthamiana* and promote mite reproduction. Plant J 86, 119–131.

Witte, H., Moreno, E., Rödelsperger, C., Kim, J., Kim, J.-S., Streit, A., Sommer, R.J., 2015. Gene inactivation using the CRISPR/Cas9 system in the nematode *Pristionchus pacificus*. Dev Genes Evol 225, 55–62.

Wybouw, N., Balabanidou, V., Ballhorn, D.J., Dermauw, W., Grbić, M., Vontas, J., Van Leeuwen, T., 2012. A horizontally transferred cyanase gene in the spider mite *Tetranychus urticae* is involved in cyanate metabolism and is differentially expressed upon host plant change. Insect Biochem Mol Biol 42, 881–889.

Wybouw, N., Dermauw, W., Tirry, L., Stevens, C., Grbic, M., Feyereisen, R., Van Leeuwen, T., 2014. A gene horizontally transferred from bacteria protects arthropods from host plant cyanide poisoning. Elife 3, e02365.

Wybouw, N., Kosterlitz, O., Kurlovs, A.H., Bajda, S., Greenhalgh, R., Snoeck, S., Bui, H., Bryon, A., Dermauw, W., Van Leeuwen, T., Clark, R.M., 2019a. Long-Term Population Studies Uncover the Genome Structure and Genetic Basis of Xenobiotic and Host Plant Adaptation in the Herbivore *Tetranychus urticae*. Genetics, doi: 10.1534/genetics.1118.301803

Wybouw, N., Kurlovs, A.H., Greenhalgh, R., Bryon, A., Kosterlitz, O., Manabe, Y., Osakabe, M., Vontas, J., Clark, R.M., Leeuwen, T.V., 2019b. Convergent evolution of cytochrome P450s underlies independent origins of keto-carotenoid pigmentation in animals. Proceedings of the Royal Society B: Biological Sciences 286, 20191039.

Wybouw, N., Van Leeuwen, T., Dermauw, W., 2018. A massive incorporation of microbial genes into the genome of Tetranychus urticae, a polyphagous arthropod herbivore. Insect Mol Biol 27, 333–351.

Wybouw, N., Zhurov, V., Martel, C., Bruinsma, K.A., Hendrickx, F., Grbić, V., Van Leeuwen, T., 2015. Adaptation of a polyphagous herbivore to a novel host plant extensively shapes the transcriptome of herbivore and host. Mol Ecol 24, 4647–4663.

Xue, W.-H., Xu, N., Yuan, X.-B., Chen, H.-H., Zhang, J.-L., Fu, S.-J., Zhang, C.-X., Xu, H.-J., 2018. CRISPR/Cas9-mediated knockout of two eye pigmentation genes in the brown planthopper, *Nilaparvata lugens* (Hemiptera: Delphacidae). Insect Biochem Mol Biol 93, 19–26.

Zhang, L., Reed, R.D., 2017. A Practical Guide to CRISPR/Cas9 Genome Editing in Lepidoptera, in: Sekimura, T., Nijhout, H.F. (Eds.), Diversity and Evolution of Butterfly Wing Patterns: An Integrative Approach. Springer Singapore, Singapore, pp. 155–172.

Zhao, C., Nabity, P.D., 2017. Phylloxerids share ancestral carotenoid biosynthesis genes of fungal origin with aphids and adelgids. PLoS One 12, e0185484.

Zhurov, V., Navarro, M., Bruinsma, K.A., Arbona, V., Estrella Santamaria, M., Cazaux, M., Wybouw, N., Osborne, E.J., Ens, C., Rioja, C., Vermeirssen, V., Rubio-Somoza, I., Krishna, P., Diaz, I., Schmid, M., Gomez-Cadenas, A., Van de Peer, Y., Grbic, M., Clark, R.M., Van Leeuwen, T., Grbic, V., 2014. Reciprocal responses in the interaction between *Arabidopsis* and the cell-content-feeding chelicerate herbivore spider mite. Plant Physiol 164, 384–399.

