## Supplementary figures and images for "Targeted mutagenesis using CRISPR-Cas9 in the chelicerate herbivore *Tetranychus urticae*"

### Figure S1

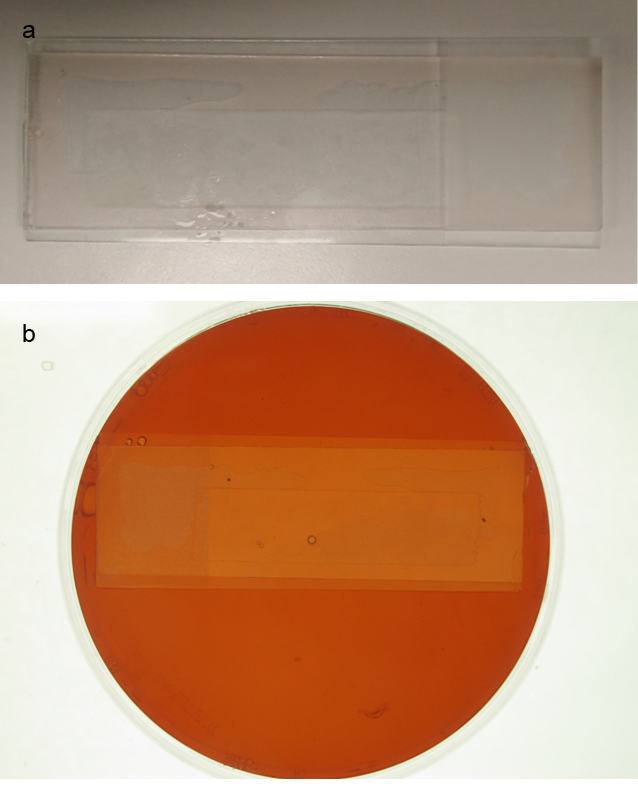

### Figure S2

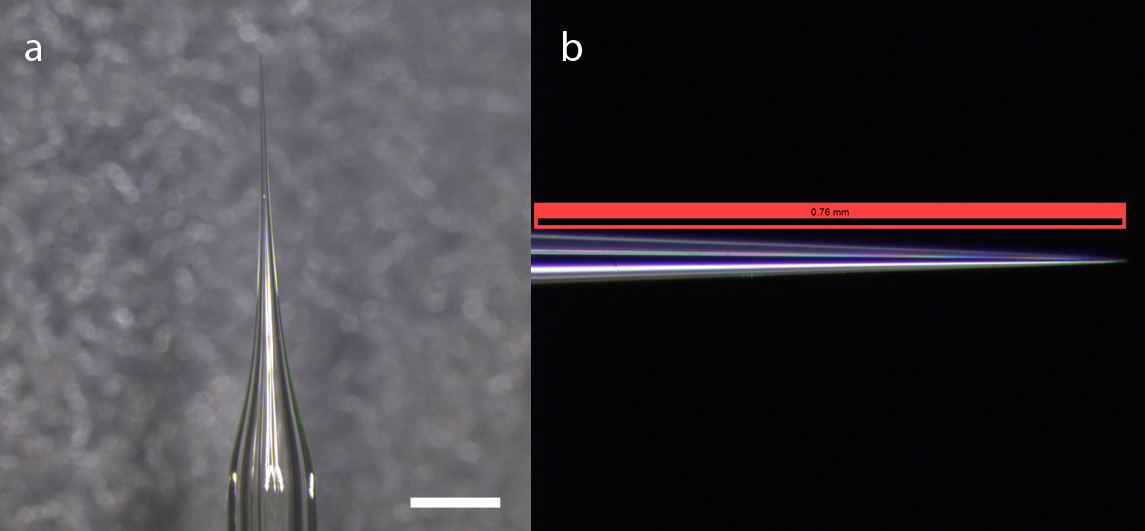

### Figure S3

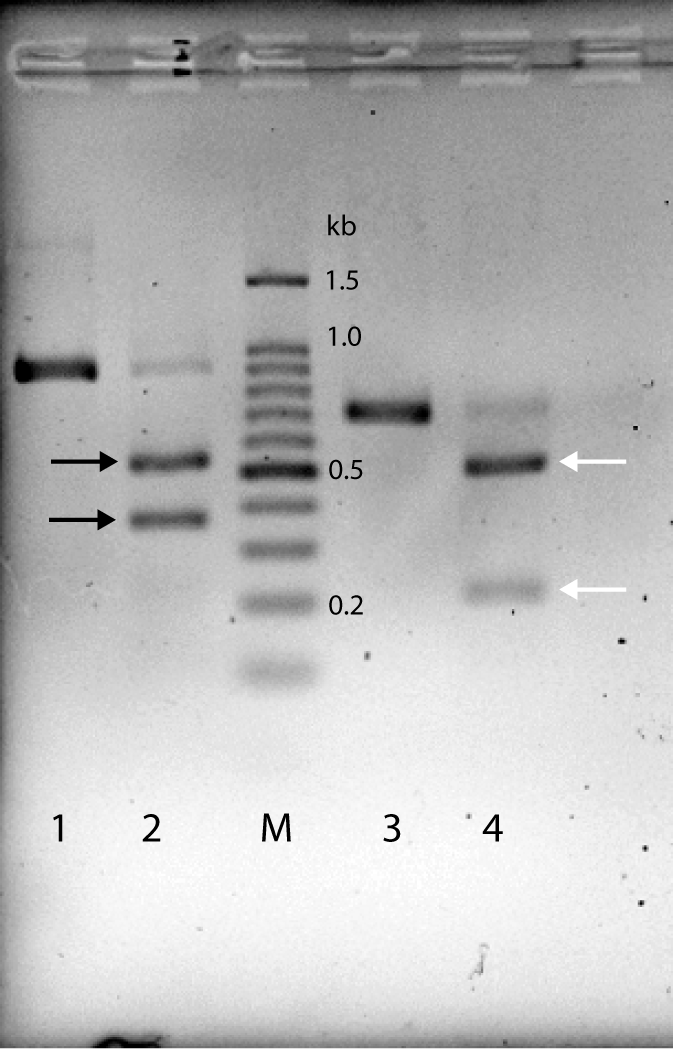

### Figure S4

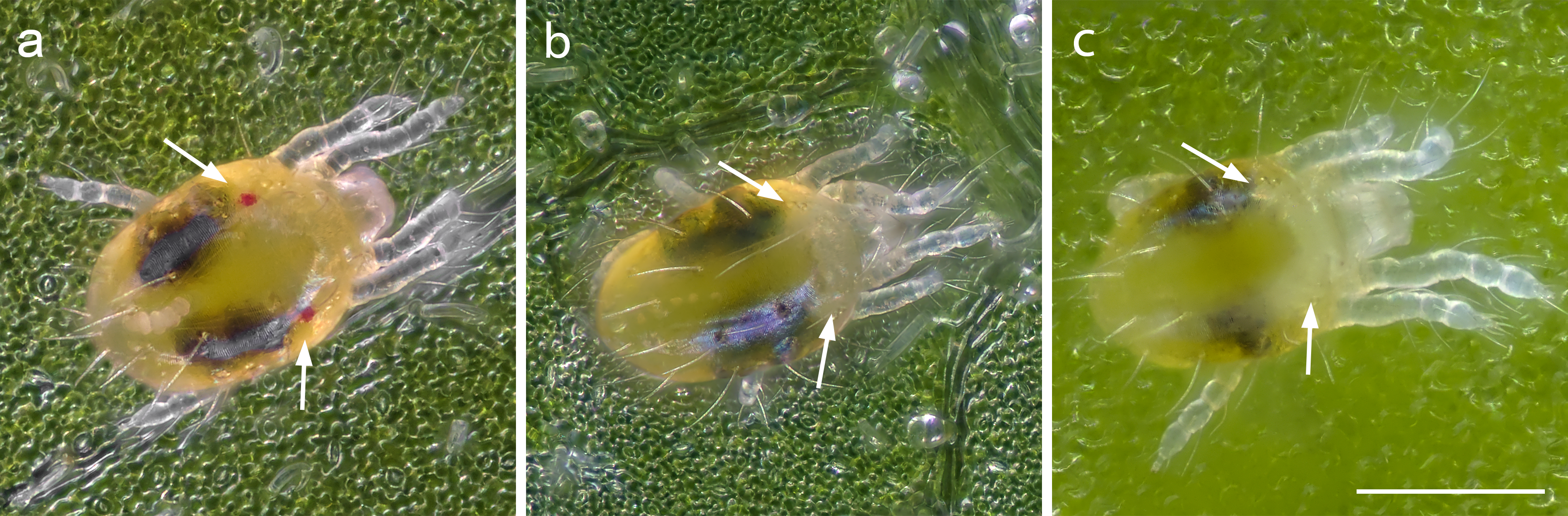

### Figure S5

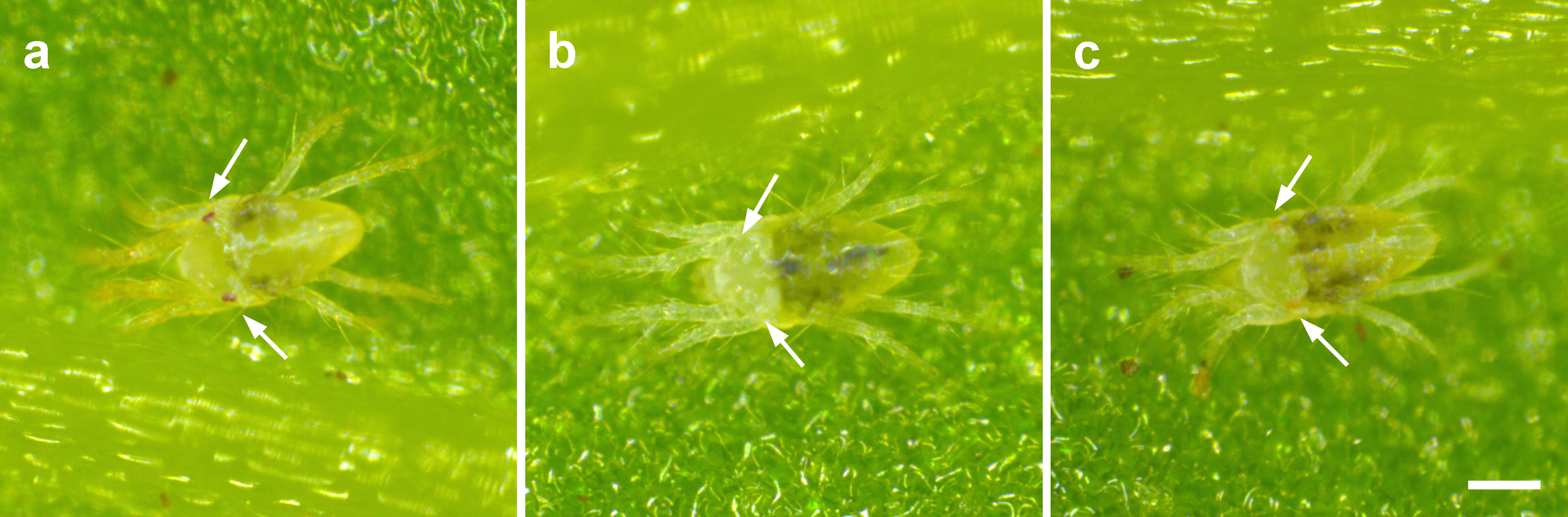
