## Supplementary material for "Targeted mutagenesis using CRISPR-Cas9 in the chelicerate herbivore *Tetranychus urticae*": Figure S6

```
tetur01g11270_WT  1    TCAGGAAAAAGAGCAATTGTGATTGGCGCTGGTGTTGGTGGTTCAGCTGT 50

tetur01g11270_WT  101  ATGATTTCAGTGGAGGCCGATGTTCGTTGATCAGAAAAAATGGACATCGA 150
tetur01g11270_A   101  ...................................C.............. 150
tetur01g11270_B   101  .................................................. 150
                                                         K->Q
tetur01g11270_WT  151  TGGGACCAAGGCCCATCACTTTACCTAATGCCTAAGCTGTTTGAGGAGAC 200
tetur01g11270_A   151  .................................................. 200
tetur01g11270_B   151  .................................................. 200

tetur01g11270_WT  201  ATTTGCTGATTTAGGTGAGGATATCAACGATCACTTGGAGCTGCTTAAAT 250
tetur01g11270_A   201  ............G..................................... 250
tetur01g11270_B   201  .................................................. 250

tetur01g11270_WT  251  GTCCGATCAATTATCGTGTTTACTTTCATGATGGCAAATTAATTGAATTG 300
tetur01g11270_A   251  .................................................. 300
tetur01g11270_B   251  .................................................. 300

tetur01g11270_WT  301  TCCAGTGATATCCAAGCAGTTTACCGACAGTTGGAAAAATTTGAGGGATC 350
tetur01g11270_A   301  .......................T.......................... 350
tetur01g11270_B   301  .................................................. 350

tetur01g11270_WT  351  AAGTGAAGATACGCTGATGAGATTTCTTGATTTTCTCAAAGAGTCTCATG 400
tetur01g11270_A   351  .................................................. 400
tetur01g11270_B   351  .................................................. 400

tetur01g11270_WT  401  TTCATTATGAGCATTCAGTTCAAATGGCTCTAAAAACACGATTCGCGTCA 450
tetur01g11270_A   401  .................................................. 450
tetur01g11270_B   401  .................................................. 450

tetur01g11270_WT  451  ATTTGGGATCTTTTCAAGTTAAAGTATATTCCTGAATTGTTTCGTATGCA 500
tetur01g11270_A   451  .................................................. 500
tetur01g11270_B   451  .................................................. 500

tetur01g11270_WT  501  TTTATACTCGACTGTTTATAAAAGAGCAACAAAGTACTTCAAAACTGAAC 550
tetur01g11270_A   501  ......................C........................... 550
tetur01g11270_B   501  .................................................. 550

tetur01g11270_WT  551  ACATGATTAAAGCTTTCACTTTCCAATCAATGTACATGGGCATGTCTCCA 600
tetur01g11270_A   551  .................................................. 600
tetur01g11270_B   551  .................................................. 600

tetur01g11270_WT  601  TATGATAGTCCTGGCCCGTACAGTTTACTTCAATACACAGAGATTGCTGA 650
tetur01g11270_A   601  .................................................. 650
tetur01g11270_B   601  .................................................. 650

tetur01g11270_WT  651  GGGTATTTGGTATCCCAAAGGTGGATTTCACAGAGTTGTCGATAAGCTGA 700
tetur01g11270_A   651  .................................................. 700
tetur01g11270_B   651  .................................................. 700

tetur01g11270_WT  701  TTGAAATTGCTTCAAATAAATTTGGCGTTAAATTTAATTACTCTGCACCA 750
tetur01g11270_A   701  .........................A........................ 750
tetur01g11270_B   701  .................................................. 750

tetur01g11270_WT  751  GTGAGGAAAATAAATGTTGATGGTAACAAAAAAGTCACTGGTATAACACT 800
tetur01g11270_A   751  ..T............................................... 800
tetur01g11270_B   751  .................................................. 800

tetur01g11270_WT  801  GGAGAGTGGAGAGGTTATTGATGCTGATTTTGTGGTGTGTAACGCTGATC 850
tetur01g11270_A   801  ................G................................. 850
tetur01g11270_B   801  .................................................. 850
                                      I->V
```

```
tetur01g11270_WT  851  TGGTATTTGCTTATAACAATTTACTTCCACCAACATCGTACGGCACTAAA 900
tetur01g11270_A   851  .............C.................................... 900
tetur01g11270_B   851  .................................................. 900

tetur01g11270_WT  901  TTGGGTTCAAAAGATCACACTTCATCTTCAATATCCTTTTATTGGGGATT 950
tetur01g11270_A   901  .................................................. 950
tetur01g11270_B   901  .................................................. 950

tetur01g11270_WT  951  GAAGGAAAAATTACCCAAATTTACTGTTCACAATGTTTTCCTGGCACAAA 1000
tetur01g11270_A   951  ........................A......................... 1000
tetur01g11270_B   951  .................................................. 1000

tetur01g11270_WT  1001 ACTATAAAGCTTCGTTCGATGAGATCTTCAAGGGACACACTCTGCCAACA 1050
tetur01g11270_A   1001 ......................A..................T.A...... 1050
tetur01g11270_B   1001 .................................................. 1050

tetur01g11270_WT  1051 CAAGCATCTTTTTATGTTAATGTGCCCAGTCGAATCGATCCAGATGCTGC 1100
tetur01g11270_A   1051 ....................C............................. 1100
tetur01g11270_B   1051 .................................................. 1100

tetur01g11270_WT  1101 TCCACCAGGTAAAGACACAATGGTTATTTTGGTACCAACAGGTTGCATGA 1150
tetur01g11270_A   1101 C.................G............................... 1150
tetur01g11270_B   1101 .................................................. 1150

                                                         sgRNA1
tetur01g11270_WT  1151 CCAATGAAAAAGGAGCAGATTTTGATGGTTTGGTGGCAAGAGCACGAGCA 1200
tetur01g11270_A   1151 ................C.............................G... 1200
tetur01g11270_B   1151 .............................................~~~~~ 1194

tetur01g11270_WT  1201 CAGGTTATTGAGACGATTGAAAAGCAAATGGGCTTTGAATCCTTTGAATC 1250
tetur01g11270_A   1201 .................................................. 1250
tetur01g11270_B   1195 ~................................................. 1244

tetur01g11270_WT  1251 TTACATTGAAACTGAAATTGTCAATGATCCAAGAACATGGAAGGAAAAGT 1300
tetur01g11270_A   1251 .................................................. 1300
tetur01g11270_B   1245 .................................................. 1294

tetur01g11270_WT  1301 TCAACCTTTGGAATGGTTCAATTCTTGGCTTAACTCATTCCATTCCTCAA 1350
tetur01g11270_A   1301 .........................G..T..................... 1350
tetur01g11270_B   1295 .................................................. 1344

tetur01g11270_WT  1351 GTTCTTTGCTTCAGGCCAAGTTTGAAGAGTCCCGTATTTGATAATTTATA 1400
tetur01g11270_A   1351 .................C.................G.............. 1400
tetur01g11270_B   1345 .................................................. 1394

                                                 sgRNA2
tetur01g11270_WT  1401 TTTTGTTGGAGCCTCGACTCAACCAGGTACTGGAGTACCCATTGTTTTAT 1450
tetur01g11270_A   1401 ...............A......~~~~~~~..................... 1443
tetur01g11270_B   1395 .................................................. 1444

tetur01g11270_WT  1451 GTGGAGCTAAATTGTTGGAAGATCAAATTGTCCAAGATAAACTTGGTAAA 1500
tetur01g11270_A   1444 .............................A.................... 1493
tetur01g11270_B   1445 .................................................. 1494
                                                    V->I
tetur01g11270_WT  1501 ACCAAGCAAACTGAAAAATTCTCGTTCGACTTTATTGGTATCTTCATTCC 1550
tetur01g11270_A   1494 .................................................. 1543
tetur01g11270_B   1495 .................................................. 1544

tetur01g11270_WT  1551 ACTCGTCCTATTATTGCTGTTTTATT 1576
tetur01g11270_A   1544 ...T...................... 1569
tetur01g11270_B   1545 .......................... 1570
```
