## Supplementary material for "Targeted mutagenesis using CRISPR-Cas9 in the chelicerate herbivore *Tetranychus urticae*": Figure S7

```
sgRNA1               1    -------------------------------------------------- 1
sgRNA2_RC            1    -------------------------------------------------- 1
tetur01g11270_Lon    1    ~~~~~~ATGAATGGTAACTCCAGTAGCTCAGGAAAAAGAGCAATTGTGAT 44
tetur11g04810_Lon    1    ~~~~~~~~~~~~~~~~~~~~~~~~.TGA.TAAG..G.AG.T...AA.C.. 26
tetur11g04820_Lon_   1    ATGAAT..T..GTC..GAAGACTCG.TGGCTCT..G.AG.T..CAA.C.. 50

sgRNA1               1    -------------------------------------------------- 1
sgRNA2_RC            1    -------------------------------------------------- 1
tetur01g11270_Lon    45   TGGCGCTGGTGTTGGTGGTTCAGCTGTTGCTGCTCGATTGGGTAAATTAG 94
tetur11g04810_Lon    27   C....G...A........ACTGA.A.CC...A..AA.C...C.C..AAT. 76
tetur11g04820_Lon_   51   ...T.G...A........ACTGA.AT.G.G....AA.C...C.C..AAG. 100

sgRNA1               1    -------------------------------------------------- 1
sgRNA2_RC            1    -------------------------------------------------- 1
tetur01g11270_Lon    95   GATTCGATGTTACTGTTTACGAGAAAAATGATTTCAGTGGAGGCCGATGT 144
tetur11g04810_Lon    77   AT.......C..C........A.....CA...A.T.C.....T...... 126
tetur11g04820_Lon_   101  AT...........CA.............CTC.GA.T.......T...... 150

sgRNA1               1    -------------------------------------------------- 1
sgRNA2_RC            1    -------------------------------------------------- 1
tetur01g11270_Lon    145  TCGTTGATCAGAAAAAATGGACATCGATGGGACCAAGGCCCATCACTTTA 194
tetur11g04810_Lon    127  ...CGA..TTAC...G...C.TT.ACC.TT..A.....AG.TATCT.G.. 176
tetur11g04820_Lon_   151  ...CGAG.TCATC..G.....T..GCC.TT..A.....AG.TATTT.G.. 200

sgRNA1               1    -------------------------------------------------- 1
sgRNA2_RC            1    -------------------------------------------------- 1
tetur01g11270_Lon    195  CCTAATGCCTAAGCTGTTTGAGGAGACATTTGCTGATTTAGGTGAGGATA 244
tetur11g04810_Lon    177  TT.GT.TGACG.TA.T.ACA..AGA.AC...AAA..GC.T..C......C 226
tetur11g04820_Lon_   201  TT.GT.TGACG.CA.T.ACA..AGA..C...AGA..GC.T.....ATC.. 250

sgRNA1               1    -------------------------------------------------- 1
sgRNA2_RC            1    -------------------------------------------------- 1
tetur01g11270_Lon    245  TCAACGATCACTTGGAGCTGCTTAAATGTCCGATCAATTATCGTGTTTAC 294
tetur11g04810_Lon    227  .GG.AA.G.GTG.TA..T.TA..G.CAA.AAAGATC..C...AA..C.TT 276
tetur11g04820_Lon_   251  .GG.T..C..TG.TA..T..A..G..CA.GAAGATC..C...AA..C.TT 300

sgRNA1               1    -------------------------------------------------- 1
sgRNA2_RC            1    -------------------------------------------------- 1
tetur01g11270_Lon    295  TTTCATGATGGCAAATTAATTGAATTGTCCAGTGATATCCAAGCAGTTTA 344
tetur11g04810_Lon    277  ..CG.A..C..G...~~~T..A.T.AC..A.AA..C..AG..TGTT.AGC 323
tetur11g04820_Lon_   301  ..CG.GA...AT...~~~T......AC..A.AC..C.GGAC.CGTT.AGC 347

sgRNA1               1    -------------------------------------------------- 1
sgRNA2_RC            1    -------------------------------------------------- 1
tetur01g11270_Lon    345  CCGACAGTTGGAAAAAATTTGAGGGATCAAGTGAAGATACGCTGATGAGAT 394
tetur11g04810_Lon    324  T...G.AA.T...CTGA.C...CCG.GCG..~~~...G.TAA.GAA.A.C 370
tetur11g04820_Lon_   348  T.A.G.AA.T..G.GGA.C..ACC..GTG..~~~A..G.CAGAGAA.AG. 394

sgRNA1               1    -------------------------------------------------- 1
sgRNA2_RC            1    -------------------------------------------------- 1
tetur01g11270_Lon    395  TTCTTGATTTTCTCAAAGAGTCTCATGTTCATTATGAGCATTCAGTTCAA 444
tetur11g04810_Lon    371  .G.AAA.....TG.GCT..AAG.A.A.CCATCG.AA..TT.GTTACCG.. 420
tetur11g04820_Lon_   395  .GGA.A.....TG...T..A.G.A.ATGCATCG.GA.ATT.GTTACCG.. 444

sgRNA1               1    -------------------------------------------------- 1
sgRNA2_RC            1    -------------------------------------------------- 1
tetur01g11270_Lon    445  ATGGCTCTAAAAACACGATTCGCGTCAATTTGGGATCTTTTCAAGTTAAA 494
tetur11g04810_Lon    421  .ACAT.T.CT.TCA.ACT.AT.ATAATG.....C.AA.GC.TTG...CTG 470
tetur11g04820_Lon_   445  .AT.T.T.CT.TCG.ACT.AT.ATA.T......C.AA.GC.TTG...CTG 494

sgRNA1               1    -------------------------------------------------- 1
sgRNA2_RC            1    -------------------------------------------------- 1
tetur01g11270_Lon    495  GTATATTCCT~~~~~~~~~~~~~~~~~~~~~~~~~~~~~~~~~~~~~~~~ 504
tetur11g04810_Lon    471  .AT...GAA.TTCTTCAACGGTATCTACCCTGCATTATTCACTAACTTTT 520
tetur11g04820_Lon_   495  CAT.GCGAA.~~~~~~~~~~~~~~~~~~~~~~~~~~~~~~~~~~~~~~~~ 504
```

```
sgRNA1              1    -------------------------------------------------- 1
sgRNA2_RC           1    -------------------------------------------------- 1
tetur01g11270_Lon   504  ~~~~~~~~~~~~~~~~~~~~~~~~~~~~~~~~~~~~~~~~~~~~~~~~~~ 504
tetur11g04810_Lon   521  ACGCTAATCTCTTCAATGATTTCATTCCTAATGTGTGGACTGGTTTGATG 570
tetur11g04820_Lon_  504  ~~~~~~~~~~~~~~~~~~~~~~~~~~~~~~~~~~~~~~~~~~~~CTAATC 510

sgRNA1              1    -------------------------------------------------- 1
sgRNA2_RC           1    -------------------------------------------------- 1
tetur01g11270_Lon   504  ~~~~~~~~~GAATTGTTTCGTATGCATTTATACTCGACTGTTTATAAAAG 545
tetur11g04810_Lon   571  ACTAATTTGA.GA.T..AT.GGCAGCCAGGAGT.AT..AA.GC.C..T.. 620
tetur11g04820_Lon_  511  ACTAATATCA.GA.C..AT.G.CAGCAC.CATG.ATT.AA.GC...GC.. 560

sgRNA1              1    -------------------------------------------------- 1
sgRNA2_RC           1    -------------------------------------------------- 1
tetur01g11270_Lon   546  AGCAACAAAGTACTTCAAAACTGAACACATGATTAAAGCTTTCACTTTCC 595
tetur11g04810_Lon   621  T.TC.AC..A.....T........CA.AG.CTGCC.....C.G..C..T. 670
tetur11g04820_Lon_  561  T.TT.ACCG...T..T....AC..CTTAG.CTGCC.....C.A..C..G. 610

sgRNA1              1    -------------------------------------------------- 1
sgRNA2_RC           1    -------------------------------------------------- 1
tetur01g11270_Lon   596  AATCAATGTACATGGGCATGTCTCCATATGATAGTCCTGGCCCGTACAGT 645
tetur11g04810_Lon   671  ....TC.T...T.T..A..A..A......AC.G..T.ATCTTTA.T...C 720
tetur11g04820_Lon_  611  ....TC.T..TT.T..A..A..A.....CAC.G..T.ATCT.TA.T.... 660

sgRNA1              1    -------------------------------------------------- 1
sgRNA2_RC           1    -------------------------------------------------- 1
tetur01g11270_Lon   646  TTACTTCAATACACAGAGATTGCTGAGGGTATTTGGTATCCCAAAGGTGG 695
tetur11g04810_Lon   721  A.TT.GGTTGCTTAT..A..A..GTCA..CC..A.A.....TG..ATG.. 770
tetur11g04820_Lon_  661  A.TT.GGTTGCTTAT..A..A..GTCA..CC..AAA.....TC..AAA.. 710

sgRNA1              1    -------------------------------------------------- 1
sgRNA2_RC           1    -------------------------------------------------- 1
tetur01g11270_Lon   696  ATTTCACAGAGTTGTCGATAAGCTGATTGAAATTGCTTCAAATAAATTTG 745
tetur11g04810_Lon   771  .A.GGCG..CA....GAC.GCTT..GAAA.T.....CGA.~~~G..AA.. 817
tetur11g04820_Lon_  711  .A.GGTG..CA.C..T...GCTT..GAA.CG.....CATT~~~..GAA.. 757

sgRNA1              1    -------------------------------------------------- 1
sgRNA2_RC           1    -------------------------------------------------- 1
tetur01g11270_Lon   746  GCGTTAAATTTAATTACTCTGCACCAGTGAGGAAAATAAATGTTGATGGT 795
tetur11g04810_Lon   818  .A..G...A.A.TAA.AAACCA.AGT..TC.C..G..TTTAA.CA.CA.A 867
tetur11g04820_Lon_  758  .A..G..GA.AC.AA.AAA.AA.GG...TGAC..G..TTTAA.A..CA.A 807

sgRNA1              1    -------------------------------------------------- 1
sgRNA2_RC           1    -------------------------------------------------- 1
tetur01g11270_Lon   796  AACAAAAAAGTCACTGGTATAACACTGGAGAGTGGAGAGGTTATTGATGC 845
tetur11g04810_Lon   868  .GTGCT.G..CA.T....G.T.A.....CCGA.AA.ACCA..G.CTTAT. 917
tetur11g04820_Lon_  808  .GT..T....CA......G.T.A......CGA...CACTA..G.CTCATG 857

sgRNA1              1    -------------------------------------------------- 1
sgRNA2_RC           1    -------------------------------------------------- 1
tetur01g11270_Lon   846  TGATTTTGTGGTGTGTAACGCTGATCTGGTATTTGCTTATAACAATTTAC 895
tetur11g04810_Lon   918  ....A....TA.T.CC.........T.AACT.A.A.......T..AC.T. 967
tetur11g04820_Lon_  858  ....A....T..T.CC..T......T.AACT.ACA.......TGC.C.T. 907

sgRNA1              1    -------------------------------------------------- 1
sgRNA2_RC           1    -------------------------------------------------- 1
tetur01g11270_Lon   896  TTCCACCAACATCGTACGGCACTAAATTGGGTTCAAAAGATCACACTTCA 945
tetur11g04810_Lon   968  .G........TAAA..T.CAGT.G.....A.CAAG.....C..TGT.C.. 1017
tetur11g04820_Lon_  908  .A..G.....TAAA..T.CAGT.G.....ACCAA......C..TGT.C.. 957

sgRNA1              1    -------------------------------------------------- 1
sgRNA2_RC           1    -------------------------------------------------- 1
tetur01g11270_Lon   946  TCTTCAATATCCTTTTATTGGGGATTGAAGGAAAAATTACCCAAATTTAC 995
tetur11g04810_Lon   1018 ..AA....TA......TC......A..G.TTCC...A.TGAAGG.C.C.A 1067
tetur11g04820_Lon_  958  ..AA.G..TA.T....TC......A..G.TTCC.G.A.TAAGGG.C.C.A 1007
```

```
sgRNA1               1    -------------------------------------------------- 1
sgRNA2_RC            1    -------------------------------------------------- 1
tetur01g11270_Lon    996  TGTTCACAATGTTTTCCTGGCACAAAACTATAAAGCTTCGTTCGATGAGA 1045
tetur11g04810_Lon    1068 AAG.......A.A...A.AC.TTCCG.T......AAGCT...T..G..T. 1117
tetur11g04820_Lon_   1008 AAG.......A.A..TA.AC.TTCCG....C.G.AAACT...T..A..T. 1057

sgRNA1               1    -------------------------------------------------- 1
sgRNA2_RC            1    -------------------------------------------------- 1
tetur01g11270_Lon    1046 TCTTCAAGGGACACACTCTGCCAACACAAGCATCTTTTTATGTTAATGTG 1095
tetur11g04810_Lon    1118 .TCA.G.CAAGA.G..AA.T..TGATG.TC....A....TCA..C.CA.T 1167
tetur11g04820_Lon_   1058 .TCA.G.CAAGA.G..AA.T..TGATG.TC....AA...TCA..C.CA.T 1107

sgRNA1               1    -------------------------------------------------- 1
sgRNA2_RC            1    -------------------------------------------------- 1
tetur01g11270_Lon    1096 CCCAGTCGAATCGATCCAGATGCTGCTCCACCA~~~~~~~~~GGTAAA~~ 1134
tetur11g04810_Lon    1168 G...A...C..G.......G.ATG.G...TAAGGATTGTATG...CGTCC 1217
tetur11g04820_Lon_   1108 G...A...C..G.........TTG.....TGAT~~~~~~~~~.....G~~ 1146

sgRNA1               1    -------------------------------------------------- 1
sgRNA2_RC            1    -------------------------------------------------- 1
tetur01g11270_Lon    1134 ~~~~~~~~~~GACACAATGGTTATTTTGGTACCAACAGGTTGCATGACCA 1174
tetur11g04810_Lon    1218 TAAGGATATG...G..C..ACCG..A.T..T..C.TC..AGAA...G..G 1267
tetur11g04820_Lon_   1146 ~~~~~~~~~~...G..C..ACCG..A.T..T..C.TC..AGAA...G..G 1186

                                                      sgRNA1
sgRNA1               1    ----------------------------....................-- 20
sgRNA2_RC            1    -------------------------------------------------- 1
tetur01g11270_Lon    1175 ATGAAAAAGGAGCAGATTTTGATGGTTTGGTGGCAAGAGCACGAGCACAG 1224
tetur11g04810_Lon    1268 G..T.C....T.GT~~~..C....A.AA...T.A..TG.TTAAGAA.A.A 1314
tetur11g04820_Lon_   1187 G.TT.CG...C.GT~~~..C..GAAAAA...T.A..A..TGAAGAA.A.. 1233

sgRNA1               20   -------------------------------------------------- 20
sgRNA2_RC            1    -------------------------------------------------- 1
tetur01g11270_Lon    1225 GTTATTGAGACGATTGAAAAGCAAATGGGCTTTGAATCCTTTGAATCTTA 1274
tetur11g04810_Lon    1315 ..A...A.A.AAC.G...G.TA..T....AA.AA.GGAT..C.CCAG.A. 1364
tetur11g04820_Lon_   1234 ..A....CA.AAT.G...C.TA..T....AA.AA.GGAT..C.CCAGCA. 1283

sgRNA1               20   -------------------------------------------------- 20
sgRNA2_RC            1    -------------------------------------------------- 1
tetur01g11270_Lon    1275 CATTGAAACTGAAATTGTCAATGAT~~~CCAAGAACATGGAAGGAAAAGT 1321
tetur11g04810_Lon    1365 G....TT..AA.C...AAA.CA..AACA...GAT..G...C.A..T..A. 1414
tetur11g04820_Lon_   1284 G....TT..A..C..ACAACCA..AACA...GATT.G...C.AA.C..A. 1333

sgRNA1               20   -------------------------------------------------- 20
sgRNA2_RC            1    -------------------------------------------------- 1
tetur01g11270_Lon    1322 TCAACCTTTGGAATGGTTCAATTCTTGGCTTAACTCATTCCATTCCTCAA 1371
tetur11g04810_Lon    1415 A...TAG....GGA..AAGC...T.A..AA.TT......GTC.A.AAA.T 1464
tetur11g04820_Lon_   1334 ....TTG....GGA..AA.C...T.A..GA.CT....CAGTT.AA.CA.T 1383

sgRNA1               20   -------------------------------------------------- 20
sgRNA2_RC            1    -------------------------------------------------- 1
tetur01g11270_Lon    1372 GTTCTTTGCTTCAGGCCAAGTTTGAAGAGTCCCGTATTTGATAATTTATA 1421
tetur11g04810_Lon    1465 T.AGCA........A..T.A.CAA...T..G.A.A......C...C.T.. 1514
tetur11g04820_Lon_   1384 T.A....A...TC.A..TCAACAA.G.T..T.A.AG...TCA...G.T.. 1433

                                                   sgRNA2
sgRNA1               20   -------------------------------------------------- 20
sgRNA2_RC            1    -------------------------....................----- 20
tetur01g11270_Lon    1422 TTTTGTTGGAGCCTCGACTCAACCAGGTACTGGAGTACCCATTGTTTTAT 1471
tetur11g04810_Lon    1515 C........T..T..A........T...G.......C..AC.A...A.G. 1564
tetur11g04820_Lon_   1434 C........T..T..A........T...G.......T..AC.A...A.G. 1483

sgRNA1               20   -------------------------------------------------- 20
sgRNA2_RC            20   -------------------------------------------------- 20
tetur01g11270_Lon    1472 GTGGAGCTAAATTGTTGGAAGATCAAATTGTCCAAGATAAACTTGGTAAA 1521
tetur11g04810_Lon    1565 .CTC....CTT.C.....TCCC.A.......AGC...CC.TTCC.C.C.. 1614
tetur11g04820_Lon_   1484 .CTC....CTT.C.....TCCC.A......AAGCC...CTCGCCAAGTC. 1533
```

```
sgRNA1             20   -------------------------------------------------- 20
sgRNA2_RC          20   -------------------------------------------------- 20
tetur01g11270_Lon  1522 ACCAAGCAAACTGAAAAATTCTCGTTCGACTTTATTGGTATCTTCATTCC 1571
tetur11g04810_Lon  1615 .AG.CT.CTG.CATCG.C.G.CAA.GA~~~~~~~~~~~~~~~~~~~~~~~ 1641
tetur11g04820_Lon_ 1534 .AGCTT.GTTTCC..TG.~~~~~~~~~~~~~~~~~~~~~~~~~~~~~~~~ 1551

sgRNA1             20   -------------------------------------------    20
sgRNA2_RC          20   ---------------------------------------------- 20
tetur01g11270_Lon  1572 ACTCGTCCTATTATTGCTGTTTTATTTTGTCTTCGGTAACAAATAA 1617
tetur11g04810_Lon  1641 ~~~~~~~~~~~~~~~~~~~~~~~~~~~~~~~~~~~~~~~~~~~~~~ 1641
tetur11g04820_Lon_ 1551 ~~~~~~~~~~~~~~~~~~~~~~~~~~~~~~~~~~~~~~~~~~~~~~ 1551
```
