## Supplementary material for "Targeted mutagenesis using CRISPR-Cas9 in the chelicerate herbivore *Tetranychus urticae*": Table S1

**Table S1 -** Composition of Cas9-sgRNA injection mix

| Component | Stock concentration | Volume in mix (μL) | Final concentration in mix (μM) |
|---|---|---|---|
| Cas9 (IDT) | 61 μM (10 mg/mL) | 2 | 29.61 |
| sgRNA 1 | 165 μM | 1.14 | 45.66 |
| sgRNA 2 | 239 μM | 0.78 | 45.25 |
| chloroquine | 10 mM | 0.2 | 485.44 |
| Total | | 4.12 | |
