## Supplementary material for "Targeted mutagenesis using CRISPR-Cas9 in the chelicerate herbivore *Tetranychus urticae*": Table S2

**Table S2** - Primers used in this study

| **Product name** | **Primer #[a]** | **Primer names & sequences (5' to 3')[b]** | **$T_m$ (°C)** | **Amplicon** |
|---|---|---|---|---|
| tetur01g11270_DNA_1 | - | F: CCCAAAGGTGGATTTCACAG | 60.34 | 895 bp |
| | - | R: GGACGAGTGGAATGAAGATACC | 59.83 | |
| tetur01g11270_DNA_2 | - | F: TGGGCTTTGAATCCTTTGAA | 60.56 | 699 bp |
| | - | R: GGGATTCTATCTTGAAAGGCAAG | 60.43 | |
| tetur01g11270_cDNA_complete | 1 | F: TGAATGGTAACTCCAGTAGCTCAG | 59.84 | 1610 bp |
| | 2 | R: GTTACCGAAGACAAAATAAAACAGC | 59.55 | |
| tetur01g11270_cDNA_intA | 4 | R: CTCTGTGAAATCCACCTTTGG | 59.58 | - |
| | 5 | F: ACGGCACTAAATTGGGTTCA | 60.37 | |
| tetur01g11270_cDNA_intB | 3 | F: TAAAAACACGATTCGCGTCA | 60.25 | - |
| | 6 | R: TCATTGGTCATGCAACCTGT | 59.97 | |

[a] see Figure 3a

[b] "F" and "R" indicate primer sequences in the forward and reverse orientation, respectively.
